# Machine learning-assisted enzyme engineering through ultra-high throughput sorting and large-scale sequence-function data generation

**DOI:** 10.1101/2025.03.30.645636

**Authors:** Jingyun Zhang, Sangeetha Shanmugam, Jing Wui Yeoh, Dan Zheng, Jan Ron Goh, Zhangyuan Lin, Chueh Loo Poh

## Abstract

Machine learning (ML) shows great promise in protein engineering but has yet to be integrated with ultra-high throughput sorting (ultra-HTS) and NGS for large-scale sequence-function data generation to harness its capability to explore wider search space and more complex mutation events. Here, we introduce PUSDA, a framework that rapidly sorts mutant libraries into multiple performance groups with good accuracy and generates large-scale sequence-function data to power ML-driven protein design. As a demonstration, PUSDA generated over five million sequence-function data of an enzyme, with data processing revealing over 1.3 million unique enzyme mutants being sorted in a single day. With a trained ML model that achieved 93.52% accuracy, we further analysed combinatorial mutation events and applied a ratio-based selection approach to design novel enzyme sequences. Validation experiment demonstrated a 16.67-fold improvement in efficiency of identifying high-performance enzymes using PUSDA compared to using ultra-HTS alone. The designed novel enzyme achieved 8.23-fold increase in productivity compared to wild type. PUSDA lays a foundation to integrate ultra-HTS, NGS, and ML for future predictive enzyme engineering, offering a data-driven tool for accelerating breakthroughs in biotechnology.

## Introduction

Directed evolution, which involves constructing and screening mutant libraries, is a pivotal method for enzyme engineering^1–4^. However, screening large mutant libraries are time-consuming and resource-intensive, driving the need for faster and more efficient approaches^1,5,6^. Recent efforts have introduced high throughput screening techniques^7–11^, including experiment automations^12,13^, fluorescence-activated cell sorting (FACS)^5,8,14–17^, and fluorescence-activated droplet sorting (FADS)^18–23^ to accelerate the screening process. However, sorting accuracy remains a challenge, necessitating additional validation of numerous candidates^24^. Deep mutational scanning (DMS) supported by next-generation sequencing (NGS) has offered a systematic approach to study sorted mutant libraries. This approach shifted blind/semi-blind laboratory-based enzyme evolution to a statistical exploration^25–35^. However, DMS has limitation to analyse multiple combinatorial mutations or uncover complex sequence variants with beneficial effects^36,37^. Moreover, DMS has difficulty to be used for designing novel protein that are not present in the NGS dataset, and it is not feasible to explore the entire sequence space even with the most advanced HTS capabilities^2,6^.

ML has recently emerged as a powerful tool in enzyme design and optimization^38–40^, particularly for handling large, high-dimensional datasets. ML excels at analysing multiple mutations simultaneously and exploring broad regions of sequence space^41^, making it possible to study and predict the effects of higher-order interactions which could be difficult to capture with traditional statistical methods^42,43^. Furthermore, ML can learn complex patterns from training dataset and provide remarkable insights into the unexplored regions of protein sequence space^5,6,38,44,45^. However, previous ML-assisted enzyme engineering studies have typically focused on limited mutation region (mainly around active sites) and small experimental dataset^41,46–55^, hampering their capability to capture the complexity of enzyme performance^6^. It is worth noting that the active sites, usually served as prior knowledge for search space design, may be unknown. Moreover, limiting the search space to active sites might overlook other regions that affect enzyme functions, such as structural stability and folding^56,57^, allosteric regulation^58^, substrate and ligand binding^59,60^, or post-translational modifications^56^. To overcome these limitations, there is need to leverage advancements in high throughput techniques, such as ultra-HTS, high throughput analysis, and NGS for large scale sequence-function data generation, and training ML models to explore wider sequence space and more complex mutation events^5,6,16,44,61^.

In this paper, we present a framework, PUSDA (Protein Mutant ultra-HTS and Data Generation for ML-assisted Enzyme Engineering) as illustrated in Figure 1. PUSDA combines i) the use of FADS based ultra-HTS for rapid sorting of large enzyme mutant libraries, ii) NGS to read the sorted mutants and generate millions of sequence-function data, and iii) ML techniques to learn and uncover key features contributing to enzyme performance, enabling the design of novel high-performance enzyme variants. To demonstrate this framework, we applied it to an important understudied flavonoid synthase enzyme (FNS I), which is involved in the production of apigenin, a flavonoid possessing multiple health benefits, including anti-inflammation, cardiovascular protection, cancer prevention, and brain health support^62,63^ (Supplementary Figure S1). Using PUSDA, a very large FNS I mutant library was sorted using FADS (enabled by an earlier developed riboswitch-based biosensor^64^) into high, medium, and low-performance groups at a rate of over one million mutants per hour and with an accuracy of 95%. The sorted mutant groups were analysed using NGS, yielding more than five million sequence-function, and processing of sequencing data revealed 1,355,459 unique mutants were sorted. The data were then filtered to enhance quality, resulting 1,119,990 sequences for ML model training (354,682 in high, 399,807 in medium, 437,501 in low-performance group). Twelve ML models were trained and evaluated, among which, random forest classifier (RFC) model using DNA sequence as input achieved the highest accuracy of 93.52%. To study more complex mutations, we analysed mutation combinations and developed a ratio-based selection approach to identify higher-dimensional features, which were then used to design novel enzyme sequences along with their predicted performance. Validation experiments demonstrated that PUSDA was 16.67 times more efficient at identifying high-performance enzymes than using Ultra-HTS alone, with the designed FNS I mutant showing 8.23-fold increase in productivity compared to the wild type. By leveraging large mutant libraries, ultra-HTS, NGS, and ML, PUSDA enables the exploration of a vast number of samples with extended search space and generates high quality, large-scale sequence-function data, which can be processed by ML to uncover complex key features, enabling more efficient and effective enzyme design. To the best of our knowledge, this is a first study that has integrated ultra-HTS with NGS to generate meaningful big sequence-function data for ML-assisted enzyme engineering. We envision that PUSDA will significantly enhance the efficiency and robustness of enzyme engineering, thereby driving advancement in the field of biomanufacturing and beyond.

**Figure 1:**
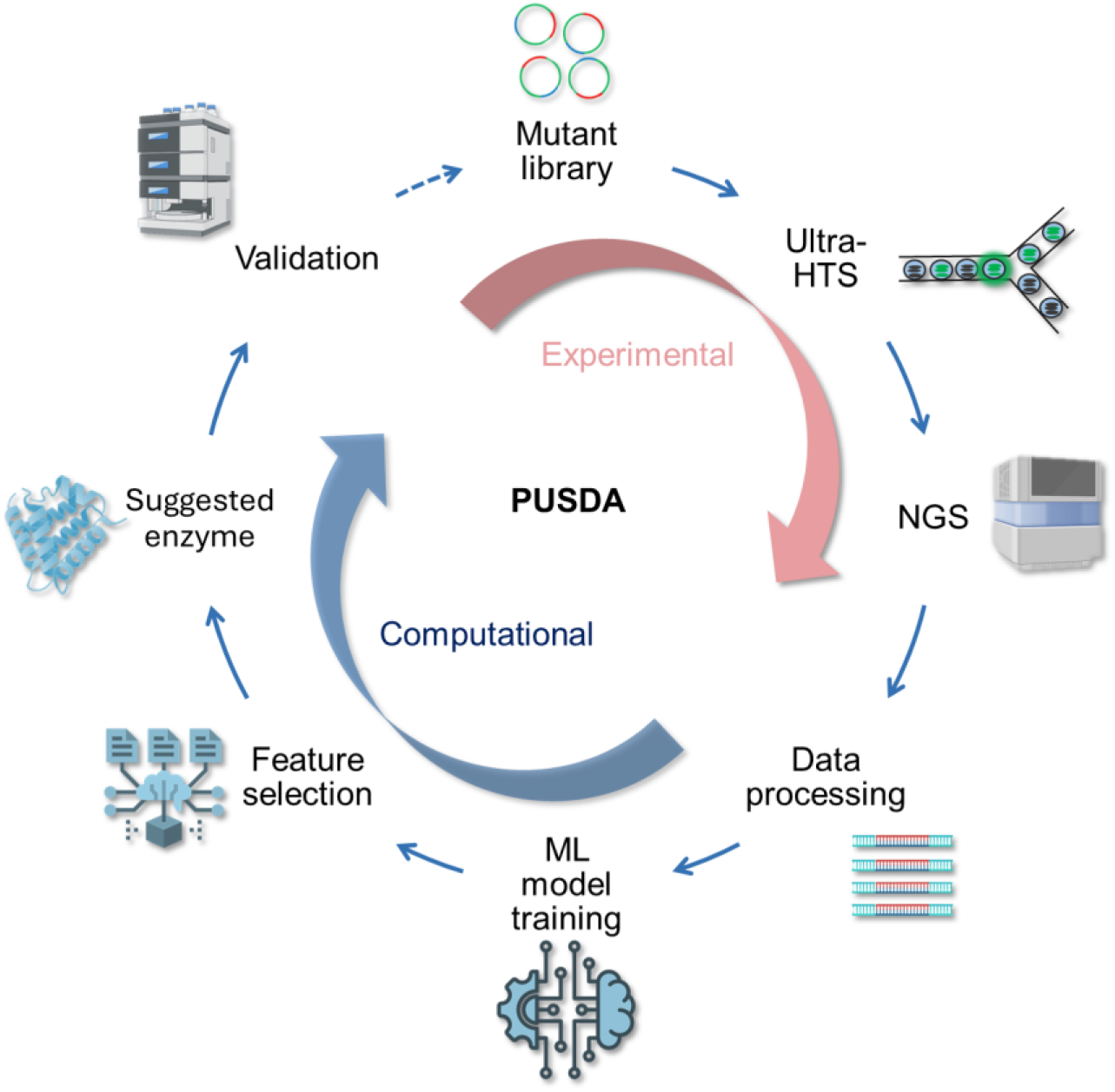
Illustration of PUSDA. The large enzyme mutant library was first constructed and sorted into different performance groups. NGS was then used to retrieve the sorted mutant sequences. The retrieved data was subsequently processed and filtered. ML models were trained using the filtered dataset for feature extraction. Higher-dimensional features were selected to design novel enzyme variants along with their predicted performance. Finally, the model suggested enzymes were experimentally validated to confirm desired performance.

## Results

### 1 Biosensor for FNS I characterization

There is growing interest in the sustainable bioproduction of flavonoids^65^, but the FNS I remains under explored, and its productivity requires further improvement. To address this, we first constructed the apigenin biosynthesis pathway in *E. coli*, in which the enzyme FNS I that converts naringenin (substrate) to apigenin (product) was regulated by an IPTG inducible promoter (Figure 2A). Directly measuring the product apigenin would enable more accurate assessment of enzyme performance and acquisition of more reliable data for training ML models. To achieve this and to facilitate high-throughput analysis/sorting, we applied an apigenin riboswitch-based biosensor developed in our previous work (Figure 2B). The biosensor produces a green fluorescence protein (GFP) readout in the presence of apigenin, otherwise remains inactive. It showed good sensitivity (Km = 6.85 µM) and an operational apigenin concentration range of 2.1 to 105 µM suitable for the analysis of the FNS I^66^. The biosensor demonstrated good specificity for apigenin, exhibiting minimal activation in response to naringenin (Figure 2B). The output level (GFP/OD_600_) when exposed to 70 µM naringenin was only 24.6% of that observed with the same concentration of apigenin. The biosensor achieved high GFP fold activation, reaching 9.1-fold at 105 µM apigenin concentration (Supplementary Figure S2). This operational range and fold activation demonstrated the capability of the biosensor to reliably detect apigenin produced by the *E. coli* in this study.

**Figure 2:**
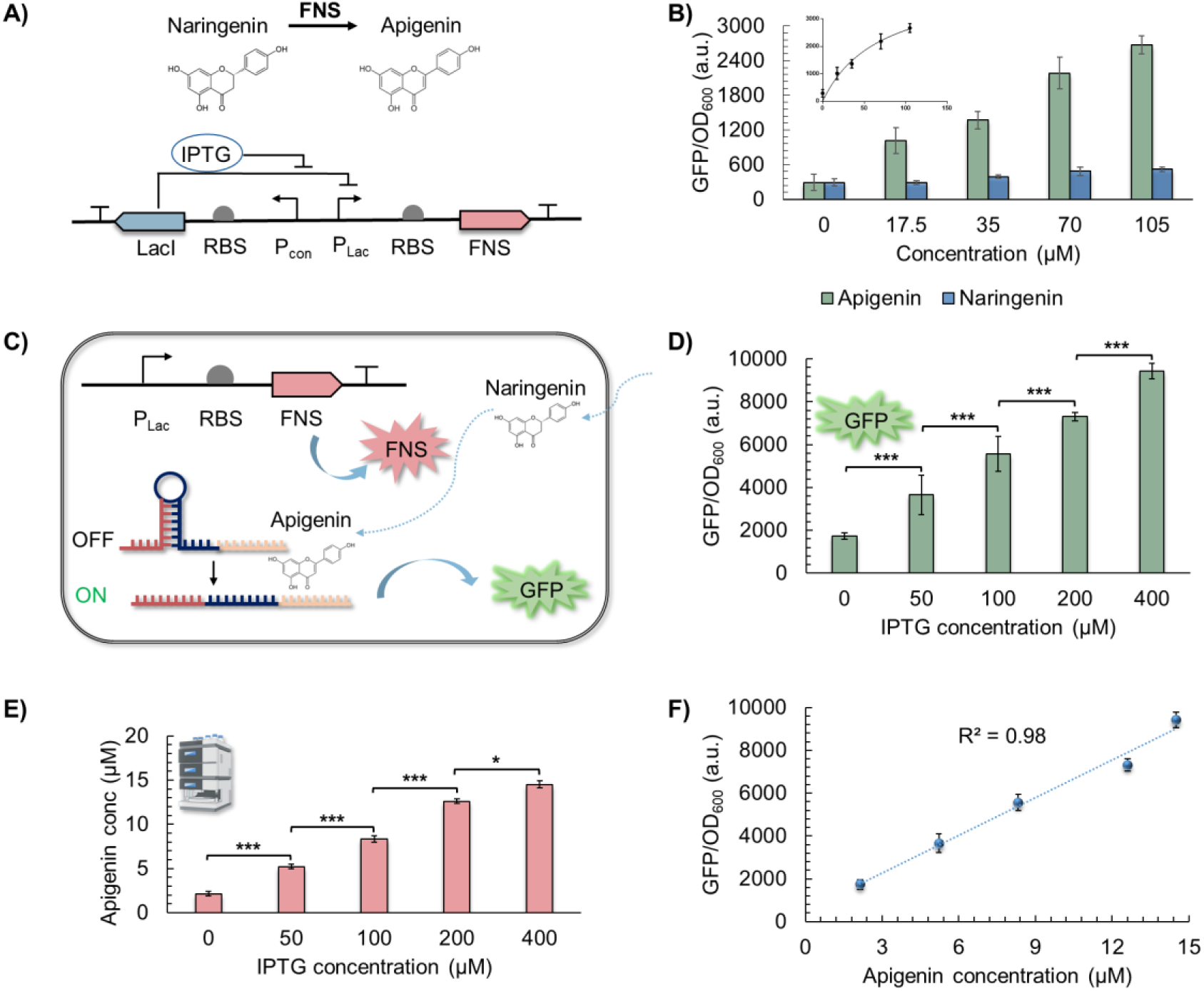
Apigenin-production pathway construction and biosensor characterization. (A) Apigenin-production pathway. FNS I was under the control of IPTG inducible promoter, upon adding IPTG, FNS I would be expressed and convert naringenin to apigenin. (B) Biosensor characterization. The biosensor expressed different GFP levels at different apigenin and naringenin concentrations. GFP were measured after 24 hours of incubation. (C) Apigenin production and *in vivo* sensing. The naringenin would first be converted to apigenin which would be then sensed by the biosensor that co-transformed into the *E. coli*. (D) Biosensor readout at different IPTG concentrations, GFP was measured after 16 hours of incubation. (E) *E. coli*-produced apigenin concentrations measured by HPLC when induced with different IPTG induction levels. (F) Correlation between biosensor readout and HPLC measurement. Data information: The experimental data were represented as mean ± S.D. (n = 3). Statistical significances of ***P < 0.001, **P < 0.01, and *P < 0.05 were calculated based on two-sample unpaired t-test.

The apigenin-producing pathway and apigenin biosensor were then co-transformed into *E. coli*. Upon adding IPTG, the cells expressed FNS, which would convert naringenin (200 µM) into apigenin. The produced apigenin would then activate the biosensor to express GFP. The apigenin production and sensing mechanism is illustrated in Figure 2C. To establish the relationship between the biosensor’s output and apigenin production, the *E. coli* strain was exposed to different IPTG concentrations. The results revealed that increasing IPTG concentrations led to higher GFP expression, reaching 5.5-fold increase at 400 µM IPTG (Figure 2D). Meanwhile, the concentrations of apigenin produced by the *E. coli* were confirmed by high performance liquid chromatograph (HPLC) measurement, where increasing IPTG concentrations also resulted in higher apigenin production, ranging from 2.1 to 14.5 µM (Figure 2E). A strong linear correlation (R^2^ = 0.98) was observed between apigenin production and biosensor readout (Figure 2F), demonstrating the biosensor’s effectiveness for *in vivo* sensing. This suggested that the biosensor holds potential for ultra-HTS of apigenin-producing mutants based on fluorescence output.

### 2 Ultra-HTS of mutant library and big data generation

Many existing studies primarily focused on modifying the active sites due to their direct impact on enzyme function^67^. However, this approach may overlook other important regions, such as dimerization sites and co-factor binding sites, which also affect the enzyme functions^68^. Expanding the search space could increase the chances of discovering higher-performance enzymes. However, this significantly increases the size of the mutant library, posing challenges when screening capabilities are limited. To explore a broader genetic diversity, we constructed a random mutagenesis library with mutations spanning 500 bp (Supplementary Table S1).

In this study, we established a workflow based on our customized FADS system (Figure 3A) which offers advantages of high-speed analysis, miniaturization, and compartmentalization, enabling efficient sorting of the large mutant library. The system achieved a sorting rate of over 1 × 10^6^ droplets per hour, with an accuracy of 96% when calibrated with constitutive GFP-expressing *E. coli* (Supplementary Figure S3). To ensure accurate sorting of the FNS I mutants, we first validated that the biosensor could detect the *E. coli*-produced apigenin within the droplet. To do this, the FNS mutant library was co-transformed with the biosensor and encapsulated into the droplets, 200 µM naringenin and 200 µM IPTG were then pico-injected into the droplets after 24 hours of incubation, and sorting was based on enzyme performance as detected by the biosensor. Post-sorting analysis revealed that mutants in the positive group (threshold of the PMT readout = 1) exhibited higher fluorescence than those in the negative group (Figure 3B). GFP distribution confirmed clear separation between the positive and negative groups (Figure 3C). To further evaluate FNS mutants’ apigenin productivity, the sorted mutants were characterized in 96-wells plate (300 µL each well). Mutants from the positive group displayed significantly higher apigenin productivity (95% producing > 20 µM) compared to the negative group (90% producing <20 µM), as shown in Figure 3D. This shows that the FADS enabled by the biosensor maintained good accuracy when sorting the FNS mutant library. Statistical analysis showed strong separation between the two groups, confirming that the biosensor’s suitability for use in the ultra-HTS system. To assess scalability, we further validated the productivity of sorted mutants in larger volume (5 mL in Falcon tube). Positive mutants maintained significantly higher productivity than negative mutants (Figure 3E, p < 0.001). This demonstrated the workflow is scalable and suitable for identifying promising candidates for larger-scale applications. Moreover, these results also showcased the system’s excellent sorting accuracy and reliability, ensuring that the data generated through our ultra-HTS is of high quality for machine learning.

**Figure 3:**
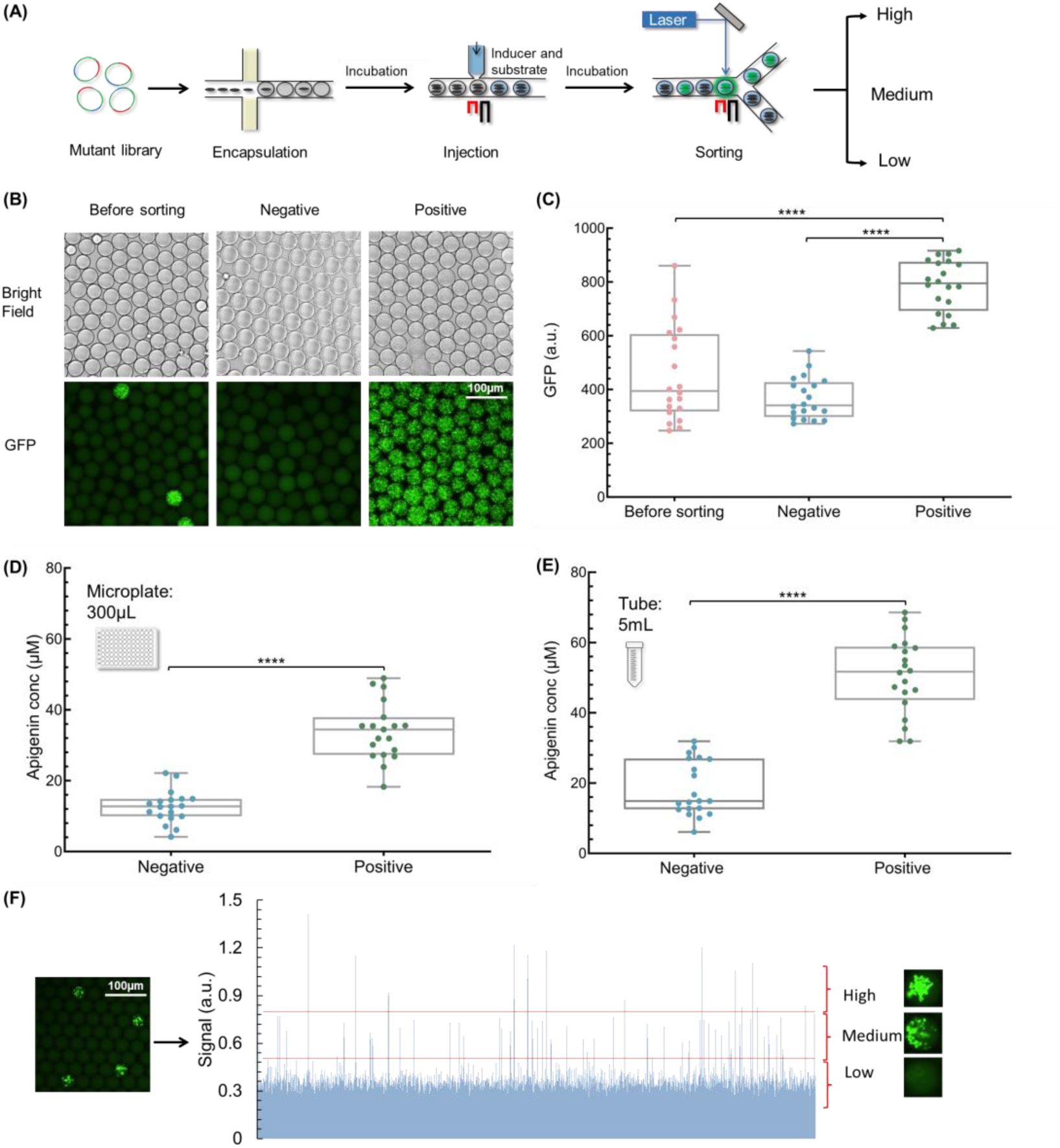
Ultra-HTS of FNS mutant library. (A) Ultra-HTS workflow. Single cell encapsulation was performed to encapsulate the single mutant into individual droplets. After 24 hours of incubation, the inducer 200 µM IPTG and substrate 200 µM naringenin were injected into the droplets. After another 24 hours of incubation, the droplets were sorted based on their fluorescence intensity. The sorter would sort the droplets with stronger fluorescence signals into the positive channel based on the set threshold. (B) Images of the droplets with encapsulated mutants before and after sorting. (C) GFP distribution of the droplets with encapsulated mutants before and after sorting. (D) Apigenin production of the sorted mutants characterized in 96-wells plate. (E) Apigenin production of the sorted mutants characterized in falcon tube. The sorted mutants remain consistent performance at larger scale. (F) Ultra-HTS of the mutant library into three performance groups based on the PMT signals. Data information: The experimental data were represented as mean ± S.D. (n = 20 per group). Statistical significances of ****P ≤ 0.0001, ***P < 0.001, **P < 0.01, and *P < 0.05 were calculated based on two-sample unpaired t-test.

With the confidence of sorting the FNS mutants using FADS and the apigenin biosensor, we conducted another round of ultra-HTS. In this round, the mutant library was sorted into three apigenin-production groups (Figure 3F): Low (L, < 6 µM), Medium (M, 6-14 µM), and High (H, > 14 µM), with PMT thresholds set at 0.5 and 0.8 to collect sufficient mutants in each group, respectively. While previous work emphasized the high-performance mutations^69^, examining mutations with varying performance levels can provide deeper insights into group-based sequence-function relationships. These datasets would be instrumental for developing, evaluating, and refining ML models.

### 3 Sequence-function data processing and analysis

To obtain sequence-function data of the sorted mutants, NGS was employed to retrieve DNA sequences from the H, M, and L groups. Over five million raw sequences were acquired from NGS. These raw sequences were first filtered to remove low-quality and redundant reads before being used for ML to identify key features. To this end, we developed an algorithmic framework to read, filter, and extract critical information (including mutant library size, mutant occurrence, unique sequences, and mutation events) from the NGS data. The data processing workflow is outlined in Figure 4A. Firstly, the sequences were filtered based on primer matching. Repeated and overlapping sequences across the three groups were removed, resulting in unique mutant libraries that consist of 423,800 sequences in H, 475,295 sequences in M, and 456,364 sequences in L group, respectively. Next, insertions and deletions that caused frameshift were removed based on BLAST alignment (Supplementary Figure S4 and S5). This resulted in the final filtered dataset of approximately 1.2 million sequences, with 354,682 sequences in H, 399,807 sequences in M, and 437,501 sequences in the L group (Figure 4B). The alignments for each sequence were used to generate a data frame, documenting the specific type and location of mutation events across the 500 bp region for each group. This created DNA sequence landscape for the H, M, and L groups (Supplementary Figure S6), with mutation event occurrences shown in Supplementary Figure S7.

**Figure 4:**
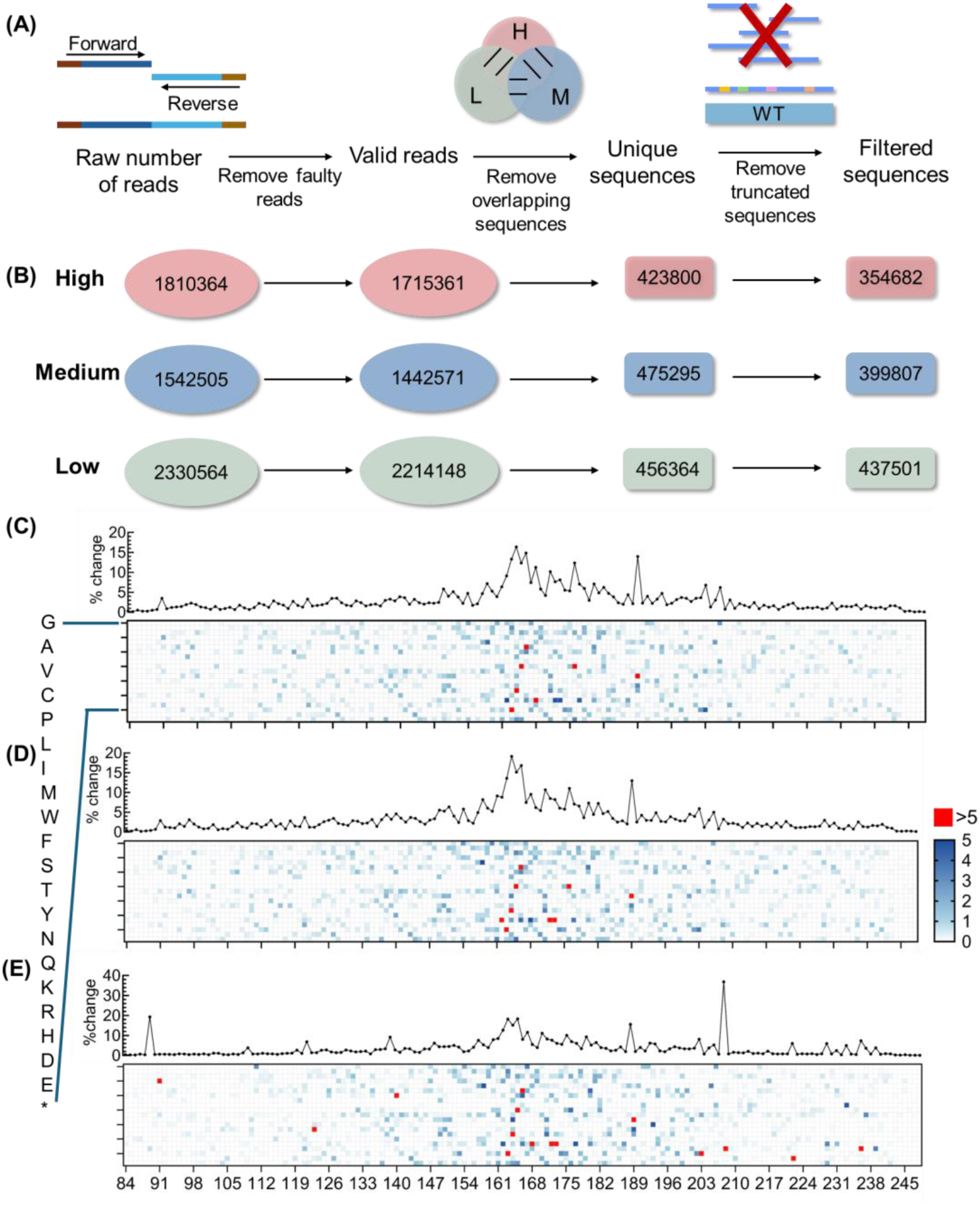
Data processing and mutant AA sequences representation. (A) Data processing workflow, including sequence construction, removal of faulty reads, identification of unique sequences, and removal of truncated reads after full length matching. (B) Number of DNA sequences in each data processing step. (C) Protein landscape of mutants in H group. (D) Protein landscape of mutants in M group. (E) Protein landscape of mutants in L group. The percentage of the change refers to the ratio of the unique sequences that have mutation on this site. 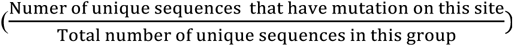

To visualise the protein landscape, the DNA sequences were converted to amino acid (AA) sequences and represented in Figure 4C-4E for H, M, and L groups, respectively. The protein landscapes revealed that most mutations occurred between AA145-AA208 in all groups. Mutations for H group clustered at AA89, AA160-AA174, AA189, AA203, and AA206. In the M group, most of the mutations were located at AA160-AA174, AA189, and AA203. In the L group, most mutations were observed at AA89, AA120, AA138, AA159-AA180, AA188, and AA205. Detailed mutation events are reported in Supplementary Figure S8, with polarity change shown in Supplementary Figure S9. In general, the L group displayed a more distinct protein landscape, while the difference between H and M were less pronounced, although the specific mutation events might be different. Since single mutation event occurrences may not be sufficient to capture higher-dimensional and more complex features that influence enzyme performance^36,37^, the unique sequences were then used for ML model training to uncover key features that differentiate the different mutant groups.

### 4 ML model training, selection, and feature extraction

Given the varying strengths and weaknesses of different ML models/algorithms, we trained several ML models to classify the three performance groups using the filtered 1.2 million sequence-function dataset (Supplementary Figure S11). The workflow for model training and feature extraction is outlined in Figure 5A. The dataset was split into 70/30 for training and testing, respectively, and sequences and mutation events were represented using one-hot encoding (Supplementary Figure S10)^70^. We trained twelve ML models, including Random Forest Classifier (RFC), Bagging Classifier (BC), Extra-trees Classifier (EC), AdaBoost Classification (AC), Gradient Tree Boosting (GTB), Histogram-Based Gradient Boosting (HGB), Decision Tree (DT), K Neighbours (KN), Stochastic Gradient Descent (SGD), Multi-Layer Perceptron (MLP) (Hidden Layers = 50), Gaussian Naïve Bayes (GNB), and Logistic Regression (LR). These models were evaluated based on precision, recall, F1-score, and accuracy (Figure 5B). Overall, most of the models achieved precision, recall, F1-score, and accuracy value above 0.8, except for GNB in L group. For the M and H groups, most models achieved scores higher than 0.6. RFC showed the best overall performance for all groups in terms of precision, recall, and F1-score, while SGD, GTB, AC, and NGB had lower scores. In terms of accuracy, RFC, EC, and MLP performed the best, achieving over 80% accuracy, while other models had varying accuracy from 38% (GNB) to 77% (HBGB) on the test dataset. GNB performed the worst, likely due to its assumption of feature independence and normal distribution^71^. Across all the models, L group was better classified than H and M groups (Supplementary Figure S12). Overall, RFC outperformed other models in handling this dataset. Hence, RFC model was selected to be used for subsequent analysis.

**Figure 5:**
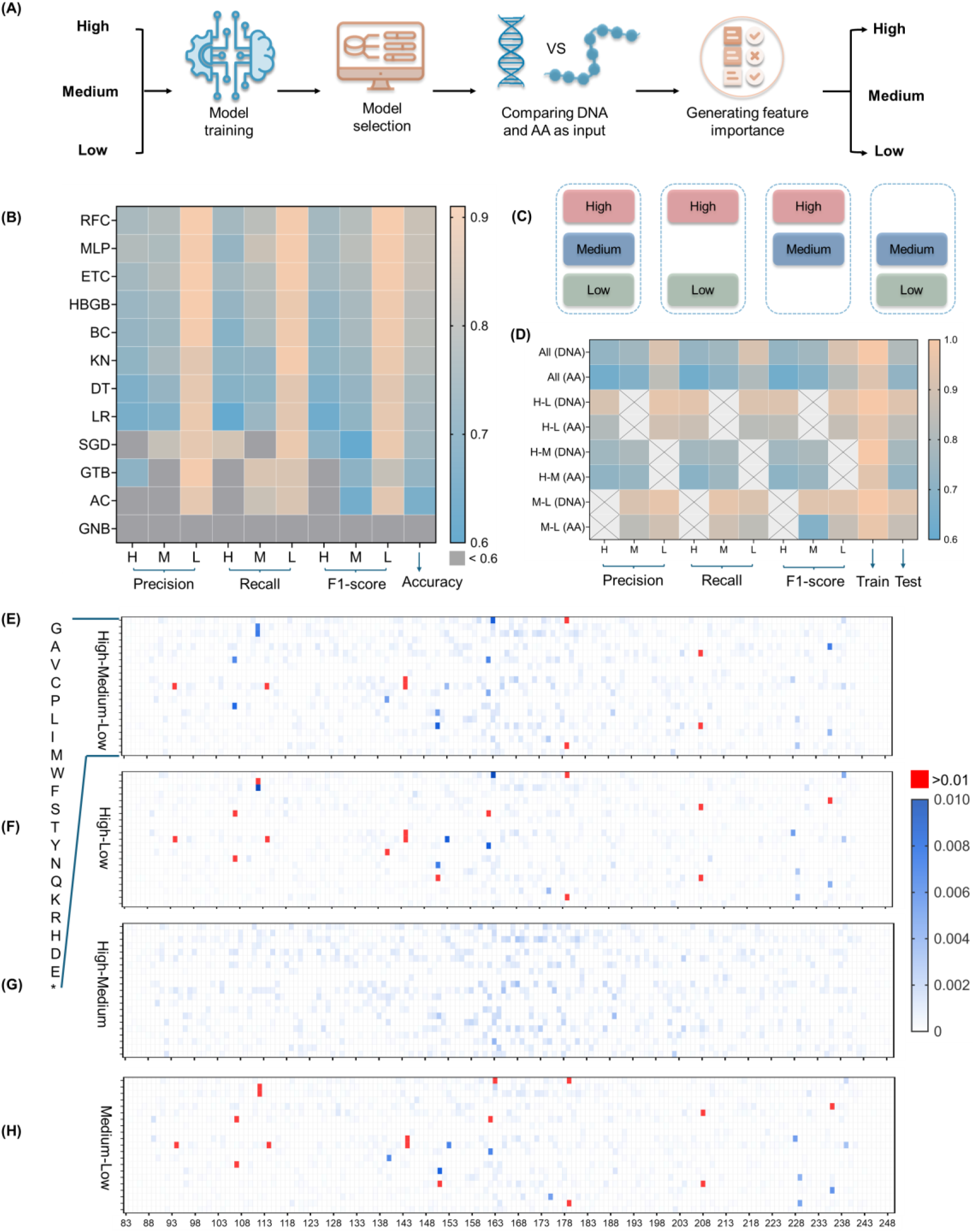
Model training, selection, and feature extraction. (A) Workflow of model training, model selection, and feature extraction. Filtered dataset was used to train several models and the models were tested using DNA or AA as input. Besides, the selected model was trained using different groups combinations. Features were then extracted and ranked to identify the important mutations. (B) Different ML models’ performance. The dataset was split into 70/30 for training and testing. (C) Running the ML model using 4 subgroups of dataset, i.e., H-M-L, H-L, H-M, and M-L. (D) Comparing the RF model performance when using DNA or AA sequence as input. (E) Feature importance scores with all three groups’ DNA sequence included as input. (F) Feature importance scores with H and L groups’ DNA sequence included as input. (G) Feature importance scores with H and M groups’ DNA sequence included as input. (H) Feature importance scores with M and L groups’ DNA sequence included as input. Red colour cells indicate a feature importance scores higher than 0.01.

In addition, as it was unclear if AA sequence would be a better input for machine learning, we evaluated the model performance using either DNA or AA sequence as input, as well as binary groups comparisons (i.e. H-L, H-M, and M-L), as illustrated in Figure 5C. The results (shown in Figure 5D) indicated that DNA sequence input outperformed AA sequence input. The accuracy of the DNA-based model was 0.81, whereas it was only 0.72 for AA input when including all three groups. For binary group comparison, the precision, recall, and F1-scores were also higher when using DNA sequences as input. The model’s highest accuracy was 0.932 for H-L and 0.935 for M-L, while the accuracy was only 0.77 when using H-M as input. This indicates that the model better distinguished L from H and M groups. More detailed comparisons of DNA and AA sequence input performance is provided in Supplementary Figure S13. Based on these findings, ML runs that used DNA sequence as input were used for subsequent ML analysis.

After model training, the features importance was assessed by evaluating the contribution of each feature to the model’s classification accuracy. DNA features were then converted to AA, and the features that did not lead to AA change were removed. The resulting feature importance landscape is shown in Figure 5E-5H. Near 2000 features were extracted for each ML run (Supplementary Figure S14). The importance score of the features in the ML runs involving the L group (H-L and M-L) were more than one order of magnitude higher (maximum 0.069) than those in the H-M comparison (maximum 0.0037). The more important features primarily distinguish the L group from the other two groups, reinforcing the need for alternative feature selection approach to capture higher-dimensional interaction between mutation events that impact the enzyme performance.

### 5 Feature selection, enzyme design, and validation

To capture more complex, higher-dimensional features, we applied feature filtration and analysed multiple mutation events in combination to identify the most important features^72,73^. The workflow for higher-dimensional feature selection is shown in Figure 6A. First, we intersected the top 30% features (Figure 6C, Figure S16) generated by the RFC model^74,75^ (feature importance dropped significantly after top 30% as shown in Supplementary Figure S15) with the mutation events that had occurrence rate of more than 1% based on BLAST analysis (Figure 6B, Supplementary Figure S17). This process resulted in a list of 504 key features (Figure 6D, Figure S18). To explore combinatorial mutations, we re-presented each mutant sequence in the database based on the presence of the 504 features, generating a new list of 669,449 mutation combinations (Figure 6E, Figure S19).

**Figure 6:**
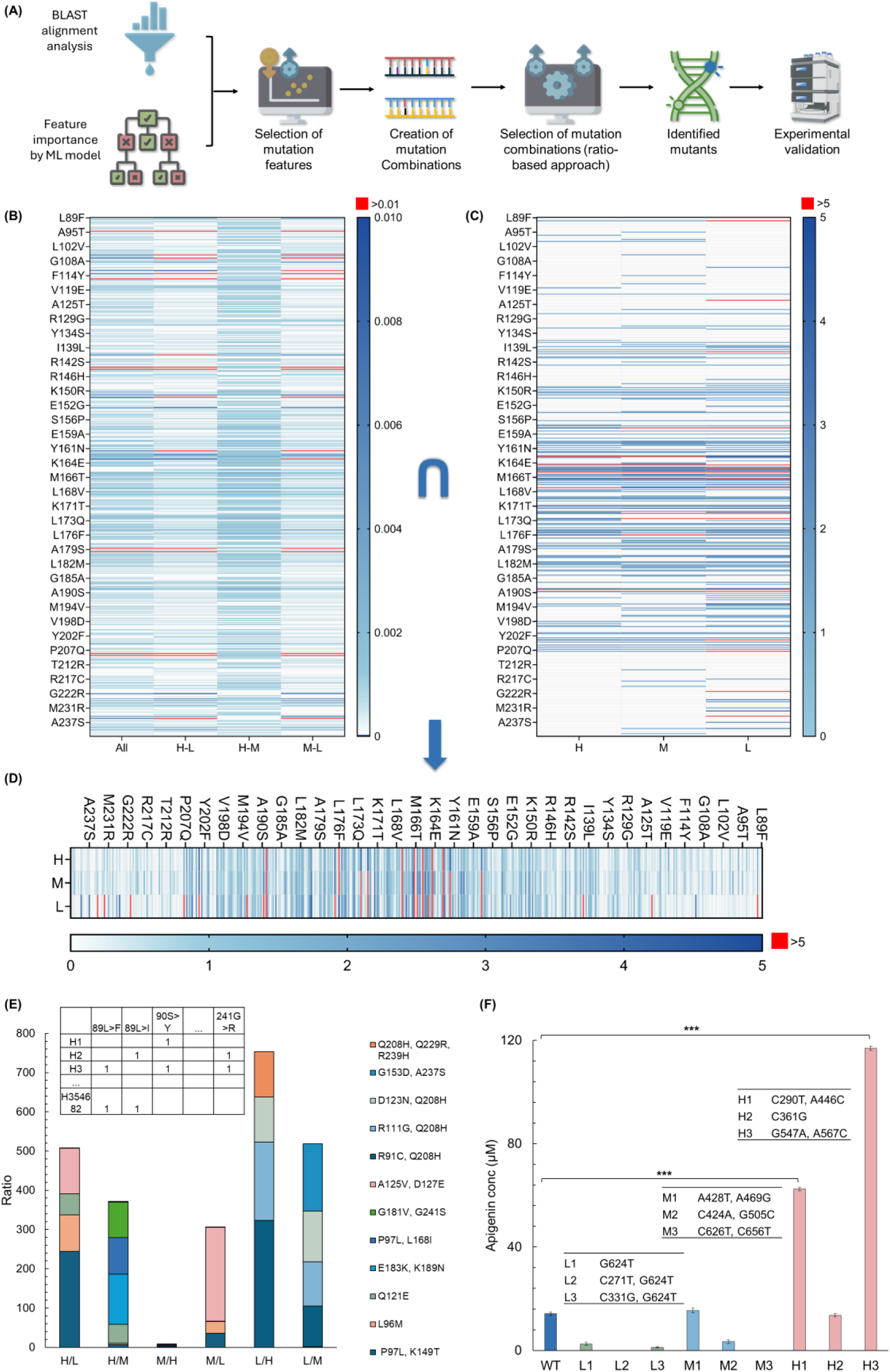
Model prediction and validation. (A) Workflow of higher dimensional feature selection and model validation. The features were filtered by combing both BLAST and ML analysis. Filtered features were then used to generate the mutation combinations. Ratio-based feature selection was subsequently used to select mutation combinations. The selected higher-dimensional features were then used to design mutant sequences. Experiments were performed to validate the designed mutants. (B) Mutation events that have occurrence more than 1% from BLAST alignment analysis. (C) Features with top 30% importance that generated by RFC model when using different group combinations as input. The feature importance generated by the ML model was used to further narrow down the mutation events that have higher importance. (D) Landscape of selected 504 key features. (E) Creation of the mutation combinations and higher-dimensional features identified from ratio-based approach. (F) Designed mutants and experimental validation. Data information: The experimental data were represented as mean ± S.D. (n = 3). Statistical significances of ***P < 0.001, **P < 0.01, and *P < 0.05 were calculated based on two-sample unpaired t-test.

Next, to identify group-associated key features from the mutation combinations, a ratio-based selection approach^75,76^ was applied, by calculating the ratio of occurrences for each mutation combinations between the groups (e.g., H/L and H/M, M/H and M/L, and L/H and L/M). This approach enabled the selection of mutation combinations with the highest group-based ratio. The most significant mutation combinations identified through this approach are shown in Figure 6E. More details of ratio-based selection approach are described in Supplementary Figure S20.

Based on this feature selection workflow, we identified nine mutants with features predicted to be the most impactful for the different performance groups. These mutants were classified as low-performance (L1, L2, L3), medium-performance (M1, M2, M3), and high-performance (H1, H2, and H3) (Figure 6F). Validation experiments were then conducted to measure the mutants’ apigenin production. The experimental results showed that all three mutants in the low-performance group produced low apigenin concentrations (<6 µM); one out of three in the medium-performance group produced medium concentrations (6-14 µM), and two out of three in the high-performance group produced high concentrations (>14 µM). The local effects of amino acid changes and the mutant’ structural analysis are shown in Supplementary Figure S21 and S22.

Notably, using PUSDA, we successfully identified novel enzyme variants (H1, H3, M1, M2, L2, L3) that were not present in the training dataset, with the best-performance novel enzyme (H3) showing 8.23-fold increase in productivity over the wild type (Figure 6F). As a comparison, a blind experiment was conducted by randomly selecting and characterizing 200 sorted mutants from the H group (Supplementary Figure S23). The highest-performing mutant from the blind experiment only showed 48% lower productivity than the model-suggested novel enzyme H3, and only 4% of the randomly selected enzymes outperformed H1, demonstrating PUSDA improved the efficiency of identifying high-performance enzyme by 16.67 times.

## Methods

### Constructing apigenin-producing plasmid

The plasmid pE6k-FNS, designed to produce luteolin from naringenin using the enzyme FNS I (UniProt, Q7XZQ8.1), was constructed using NEBuilder HiFi DNA Assembly Master Mix (New England Biolabs, E2621). The DNA sequence for FNS I was synthesized by Integrated DNA Technologies (IDT, USA). This gene was inserted downstream of an IPTG-inducible promoter in the pBbE6k-RFP plasmid vector, replacing the RFP gene. The pBbE6k-RFP plasmid was a gift from Jay Keasling (Addgene plasmid # 35288; http://n2t.net/addgene:35288; RRID: Addgene_35288 ^77^). The assembled plasmids were then transformed into Acella *E. coli* via heat shock and cultured in Luria Broth (Invitrogen, 12795-027) media for recovery.

### Characterization of dose-response of the apigenin biosensor

Cells were inoculated from a glycerol stock and incubated overnight in Luria Broth (Invitrogen, 12795-027) with 25 μg/ml chloramphenicol (Sigma Life Science, C0378; diluted in 100% ethanol, Sigma-Aldrich, E7023) at 37°C with shaking at 225 rpm. The next day, 50 μL of the overnight culture was transferred into 5 mL of fresh LB containing 25 μg/ml chloramphenicol and incubated for 2 hours. After measuring the optical density at 600 nm (OD600), the culture was diluted to an OD600 of 0.1 using fresh LB with 25 μg/ml chloramphenicol. Naringenin and apigenin were added respectively to the cultures at final concentrations of 17.5, 35, 70, and 105 μM. These mixtures were transferred to a 96-well plate (Greiner Bio-One, 655180), with 300 μL per well. Control wells contained cell mixture only, without flavonoids. Each sample was prepared in triplicate. Cell growth (measured by absorbance at 600 nm) and biosensor response (quantified by GFP fluorescence with excitation at 470 nm and emission at 520 nm, using a gain of 75) were monitored using a microplate reader (Cytation 5, Biotek). The plate was kinetically shaken, and readings were taken every 10 minutes over 24 hours. All data were baseline-corrected using the readings from LB with 25 μg/ml chloramphenicol.

### *In-vivo* sensing of apigenin

The apigenin-producing plasmid pE6K-FNS and the apigenin biosensor were co-transformed into *E. coli* Acella. Cells were inoculated from a glycerol stock and cultured overnight in LB medium supplemented with 50 μg/ml kanamycin and 25 μg/ml chloramphenicol at 37 °C with shaking at 225 rpm. The following day, 50 μL of the overnight culture was transferred into 5 mL of fresh LB and incubated for 2 hours. After measuring the cell density, the culture was diluted to an OD600 of 0.1 using fresh LB. A range of IPTG concentrations (0, 50, 100, 200, 400 µM) was added, while the substrate naringenin was kept constant at 200 µM. The mixtures were dispensed into a 96-well plate (Greiner Bio-One, 655180) with 300 μL per well, with each sample prepared in triplicate. Cell growth (OD_600_) and GFP fluorescence (excitation at 470 nm, emission at 525 nm) were measured over a 24-hour period at 30°C using the microplate reader. At the experiment’s endpoint, the cell cultures from the 96-well plate were collected and analysed using HPLC (Hitachi; Pump: CM5110; Autosampler: CM5210; Column Oven: CM5310; Diode Array Detector: CM5430)to quantify the concentration of apigenin produced. The correlation between GFP fluorescence and the apigenin concentration produced by *E. coli* measured by HPLC was then analyzed.

### Construction of Mutant Library

To generate the mutant library, error-prone PCR^78^ was employed. Mutant FNS gene fragments were amplified using primers (Supplementary Table 1) flanking the wild-type FNS gene, using GeneMorph II (Agilent, 200550). The resulting amplicons were visualized on a 1% agarose gel, cast in TAE buffer, pre-stained with SYBR Safe (Invitrogen, 2265984), and gel-extracted using the QIAquick Gel Extraction Kit (Qiagen, 28706). The purified amplicons were assembled with the NEBuilder HiFi DNA Assembly Master Mix (New England Biolabs, E2621). The assembled product was transformed into *E. coli* Acella chemically competent cells, using a heat-shock method. Successfully transformed clones were picked and cultured overnight (>12 hours) at 37 °C in a shaking incubator at 225 rpm. A portion of each culture was used to prepare cryostocks (25% glycerol), while the remainder was used for downstream characterization and plasmid extraction (Qiagen, QIAprep Spin Miniprep Kit, 27106). Extracted plasmids were sent for Sanger sequencing (1st Base, Singapore) to verify the induced mutations.

### Characterization of apigenin-producing strain in falcon Tubes

50 mL conical-bottom tubes (Greiner Bio-One, 227261) containing 5 mL of LB Broth (Invitrogen, 12795-027) with appropriate antibiotics, i.e., 25 μg/mL chloramphenicol and 50 μg/mL kanamycin (Sigma Life Science, 60615; diluted in nuclease-free water) were inoculated from glycerol stock cultures and grown overnight (> 12 hours) at 37 °C in a shaking incubator at 225 rpm. The following day, at time T = 0 hours, 50 μL of this culture was diluted 1:100 into a fresh 50 mL conical tube containing 5 mL of LB media, the same antibiotics, and inducer (200 μM IPTG (Sigma Aldrich, I6758) and 200 μM naringenin (Sigma-Aldrich, N5893). The cultures were incubated at 30 °C with shaking at 225 rpm. At 24 and 48 hours, 850 μL aliquots were taken from each sample to analyse naringenin concentrations and apigenin yield using HPLC. Additionally, 30 μL aliquots were collected at these time points to measure fluorescence and OD_600_ using a microplate reader (Cytation5, BioTek, USA) with the following settings: gain of 75, excitation at 470 nm, emission at 520 nm, and OD_600_ at 600 nm.

### HPLC for flavonoids measurement

Unless specified otherwise, extracted aliquots from each sample were spun down at 13.5k rpm (17.5k g) for 2 minutes to pellet the cells. 750 μL of the supernatant was mixed with an equal volume (750 μL) of ethanol (Sigma-Aldrich, E7023) to increase the solubility of naringenin and apigenin. The mixture was syringe filtered with a 0.2 μm PES membrane (PALL Corporation, Acrodisc 4612) into a 2 mL HPLC glass vial (Supelco, 29379-U). Detection of naringenin and apigenin was performed using a Chromaster CM5000 HPLC (Hitachi; Pump: CM5110; Autosampler: CM5210; Column Oven: CM5310; Diode Array Detector: CM5430). The column temperature was set to 40°C, with a flow rate of 1.5 mL/min and an injection volume of 10 μL. Analytes were separated on a LaChrom C18-AQ 5 μm (Hitachi) HPLC column using 40% methanol (Sigma-Aldrich, 34860) and 0.6% acetic acid (Sigma-Aldrich, A6283) as mobile phases over 30 minutes. UV absorbance was detected at 300 nm with a sampling frequency of 2.5 Hz. Data was collected and processed using Chromaster System Manager Version 1.2. The amounts of naringenin and apigenin were quantified by measuring peak areas at 13 minutes and 28 minutes, respectively, and concentrations were determined from a calibration curve.

### Microfluidic device fabrication

Three types of microfluidic devices (i.e., droplet generator, pico-injector and droplet sorter) fabricated from poly (dimethylsiloxane) (PDMS, Sylgard 184™, Dow Corning Inc, Midland, MI) through soft lithography techniques ^79^ were used in the study. Single cell encapsulation process was performed using a droplet generator (40 µm width, 30 μm height, Supplementary S3). Pico-injection process was conducted using a pico-injector (25 µm width, 25 μm height, 10 μm injector nozzle, Supplementary Information S3). To sort droplets, a droplet sorter (30 µm width, 30 μm height, Supplementary Information S3) modified from Mazutis *et al.* ^80^ was used. Glass slides coated with a thin PDMS layer were used as the substrates for droplet generators, whereas uncoated glass slides were used as the substrates for pico-injectors and droplet sorters. The PDMS replicas of the microfluidic devices were securely bound to the substrates using a plasma cleaner (PDC-32G, Harrick Plasma, Ithaca, NY). Post to plasma bonding, to create the electrodes of pico-injectors and droplet sorters, the devices were heated to 80 °C, allowing the low-melting point indium alloy wire (WIREBN-52189, Indium Corporation, Clinton, NY) to melt and flow into the electrode inlet channels. Subsequently, the pico-injectors and sorters were assembled by connecting wires to these metal electrodes, allowing them to receive AC electrical signals for operation.

### Droplet encapsulation

Flow control for both aqueous and oil phases in all processes was managed using syringe pumps (Harvard Apparatus, PhD 2000). The oil phase consisted of Novec™ HFE-7500 oil (3M, Singapore) with 0.25% (w/w) Picosurf-1™ surfactant (Sphere Fluidics, Cambridge, UK). To generate single cell encapsulated droplets, the mutant libraries were diluted to approximately 8.0 × 10^6 cells/mL in LB medium with antibiotics. Two streams of the diluted *E. coli* cell suspension (flow rate: 2 μL/min) and a stream of oil (flow rate: 10.5 μL/min) were co-injected into a droplet generator.

### Droplet pico-injection

After the droplets were incubated for 24 hours, they were pico-injected with naringenin and IPTG solution. Flow rates of injector, spacing oil, and droplet maintained at about 0.25, 1.2, and 0.7 µL/min for stable and consistent injections. To disrupt the interface of the droplets for pico-injection, a 2 Vpp 20 kHz sinusoidal wave was amplified 100-fold and applied to the electrodes of a pico-injector. The concentration of naringenin and IPTG solution in each droplet was approximately 200 µM. Thereafter, the droplets were incubated at 30 °C for additional 24 hours.

### Microfluidic sorting

The optical setup for droplet sorting included a fluorescence light source (CoolLED pE-4000) and a customized photomultiplier tube (PMT) detection system. Droplet and oil flow rates were set at 0.5 μL/min and 15 μL/min, respectively. Voltage signals from the PMT were split for simultaneous signal recording and sorting. PMT analog signals were converted to digital via a data acquisition card (USB-6002, National Instruments, USA) and recorded using MATLAB’s Analog Input Recorder (MathWorks R2021a, USA). For sorting, PMT signals were processed in real-time by an Arduino DUE microprocessor (Arduino, USA), which controlled an Arduino UNO microprocessor (Arduino, USA) to produce 8 Vpp, 10 kHz square waves. These signals were amplified 100-fold by a high-voltage amplifier (Trek, model 2210) and directed to the sorter’s electrodes. When a PMT signal exceeded the threshold, sorting waves directed the droplet to the collection channel. The sorted droplets were collected from the microfluidic chip for DNA extraction (QIAprep, QIAGEN, German). The DNA were sent for NGS (PE250, NovogeneAIT, Singapore)

### Data processing and filtration

The NGS sequencing data, consisting of 500 bp reads, was divided into two halves, each flagged by either the forward or reverse primer. Constructs with correctly matched forward and reverse primers were identified, with the second half reverse complemented for final stitching. Nucleotide sequence comparisons were performed using the BLAST ^81^. Query and subject sequences were analysed for similarity and alignment scores, and the results were saved in XML format. The XML files were parsed using the Biopython NCBIXML module and custom scripts to extract information including the matching regions of the mutant sequence with the wild-type (WT) sequence, the number of gaps in the alignment, and individual mismatch positions. Read statistics, match lengths, and alignment details were stored in the Pandas DataFrame. Samples were filtered to include only full-length matches with substitution events (no gaps), resulting in a filtered dataset containing 354,682 sequences in the H group, 399,807 in the M group, and 437,501 in the L group. BLAST was used to align the matches for each sample, generating a DataFrame that captured the location and type of substitution events across the 500 bp for each performance group in the filtered dataset.

### ML model training

A total of twelve machine learning models were tested using the default hyperparameters from the scikit-learn package^82^. The dataset was split 70/30 for training and testing. One-hot encoding was used to represent the sequences^83^, indicating the presence or absence of a nucleotide at each position. The output of the models was categorical, ranking the sequences into three functional groups (Low, Medium, High). The models tested from scikit-learn included: Bagging Classifier, Random Forest Classifier, Extra-Trees Classifier, AdaBoost Classifier, Gradient Tree Boosting, Histogram-based Gradient Tree Boosting, Decision Tree, K-Nearest Neighbors, Stochastic Gradient Descent, Multi-Layer Perceptron, Gaussian Naïve Bayes, and Logistic Regression. Model performance was evaluated by reporting precision, recall, and F1-score for each classification group, as well as overall accuracy. The Random Forest model was selected for further analysis. Both DNA and amino acid sequences, represented as one-hot encoded inputs, were tested with a 70/30 train/test split while keeping the same categorical output.

### Studying mutation combinations

The top 30% of features, based on feature importance rankings from four independent Random Forest runs, were selected after excluding unchanged bases compared to the wild-type sequence. These features were then combined with mutation events present in more than 1% of sequences in each of the three performance groups (Low, Medium, High), as identified through BLAST analysis, to generate a list of DNA base mutations. Mutations that did not result in amino acid changes were excluded, resulting in a final list of 504 mutation events. Each sample from the filtered dataset was represented by the presence or absence of these mutations, creating a mutation profile for every sample across all three performance groups. This representation was similar to the one-hot encoding method used previously. The frequency of different mutation combinations was then calculated, and the combinations were sorted by their occurrence percentages across the dataset.

### Ratio-based feature selection and mutant sequence suggestion

The mutation combinations were categorized into three groups: (i) those common to all three performance groups, (ii) those shared between each pair of groups (e.g., between H and L), and (iii) those unique to each individual group. For each category, the mutation combinations relevant to each performance group were identified by calculating the ratio of occurrences (number of times a mutation combination occur) between groups, i.e. H/L, H/M, M/H, M/L, L/H, and L/M (Figure 6E). The combinations with the highest ratios for each performance group were selected, leading to the identification of mutations H1, H2, H3 for the High group, M1, M2, M3 for the Medium group, and L1, L2, L3 for the Low group for further experimental testing.

## Discussion

This paper introduces PUSDA, a framework for ML-assisted enzyme engineering through ultra-HTS and large-scale data generation. As a demonstration, PUSDA was applied to improve an enzyme FNS I. PUSDA first employed a FADS-based ultra-HTS system to sort a large mutant library (spanning a 500 bp mutation region) into three bins based on their performance in apigenin production, achieving a sorting rate over million mutants per hour with 95% accuracy. Scalability tests confirmed that mutants sorted in 10^-12^ µL droplets maintained consistent performance in larger volumes (e.g., 300 µL in microplate and 5 mL in Falcon tube). This validates that the robustness of the sorting methods across various volumes, and ensures the reliability of the generated data. Coupled with NGS and data processing, PUSDA generated more than five million sequence-function data in a single run. To enhance the data quality, a data filtering algorithm was implemented to retain only unique sequences for ML training. Twelve ML models were trained and evaluated, with the best achieving 93.52% accuracy using DNA sequence as input. Higher-dimensional mutation interactions were also studied, and a ratio-based feature selection approach was developed to identify group-specific characteristics. Importantly, PUSDA could be used to design novel enzyme variants, and experimental validation revealed 16.67-fold improvement in the efficiency of identifying high-performance enzymes compared to using ultra-HTS alone. One of the novel enzymes FNS I variant, absent from the training dataset, showed 8.23-fold increase in productivity. These findings exemplify PUSDA’s capability to design novel improved enzymes, while greatly reduced reliance on laborious trial-and-error experiments.

Conventional enzyme screening methods are constrained by throughput limitation^84,85^, and while recent advances in HTS techniques have improved the mutant library analysis speed^12,14,86^, assay limitation remain a bottleneck^6^. Genetically encoded biosensors offer a promising solution by enabling higher throughput^54^. However, some studies have largely relied on indirect sensing (e.g., NADH biosensor), which do not always accurately correlate with enzyme catalytic performance^87–90^. By contrast, our work employs a biosensor that directly detects the target product, allowing more precise evaluation of the enzyme performance and generating higher-quality data for ML applications^5^. This approach demonstrates PUSDA’s potential to employ other biosensors^91,92^ for diverse enzyme development projects.

Previous ML-assisted enzyme engineering efforts have often been constrained by small dataset and narrow search space (primarily on enzyme active sites)^41,46–55^. This limits their ability to capture more complex mutations and wider mutation regions which has potential to improve enzyme performance^6^. PUSDA addressed these challenges by leveraging the advantages of ultra-HTS, NGS, and ML to investigate expanded search space, sort a larger mutant library, generate larger dataset, and reduce reliance on prior knowledge of enzyme structure and active sites. Unlike prior methods which focused on screening for high-performance mutants alone^30,33–35^, this work sorts mutants into high, medium, and low-performance groups, and analysed all groups in different combinations. This created a richer dataset, enabling the ML models to gain deeper insights and uncover more nuanced sequence-function relationships. Instead of using traditional statistical analysis methods^25–34^, PUSDA employs ML to handle larger dataset with complex mutation events, and design novel mutant sequences based on enzymes performance prediction. ML models vary in their assumptions and strengths (e.g., GNB assumes feature independence and a Gaussian distribution^93^), PUSDA evaluated 12 models and explored different input formats (DNA vs. AA) to identify the best fit. The models showed different levels of accuracy, with RFC model achieved the highest accuracy at 93.52%. This highlights the importance of testing different ML models for optimal performance. Besides, this work introduced a ratio-based selection method to analyse complex mutation relationships, moving beyond the traditional focus on single mutation events. This approach facilitates the design of mutants with multiple mutations, offering a more comprehensive strategy for directed evolution.

While the study showed that there is good predictive accuracy by the ML between high and low-performance mutants, the prediction between high and medium-performance groups was less accurate, likely due to less distinct sequence differences between these groups. Nonetheless, the high and low-performance mutants designed by ratio-based selection approach achieved good predictive accuracy, demonstrating the framework’s effectiveness in prioritizing high and low-performance variants.

This work is the first to leverage advances in ultra-high throughput experimental techniques (ultra-HTS, NGS) and machine learning to efficiently explore broader mutation region and uncover complex sequence-function relationships for enzyme engineering. Since ML models together with its training data typically need to be tailored for different protein families^94^, PUSDA addressed an important gap in efficient and intelligent enzyme engineering by rapidly generating the large-scale dataset. It lays the foundation for broader applications, including protein, pathway, and strain engineering. Furthermore, this study holds potential to complement advanced ML models that rely on data from existing protein databases, such as AlphaFold^95^, large language models^96,29^, and neural network-based model^45^. As ultra-HTS technologies, NGS, and computational power/methods such as machine learning/AI continue to advance, the integration/convergence of these technologies will unlock new possibilities in biology, medicine, and biomanufacturing.

## Data availability

Data will be made available on request.

## Code availability

Code used will be made available on request.

## Acknowledgements

The authors gratefully acknowledge the funding provided by the Singapore National Research Foundation (NRF) under the NRF Synthetic Biology Program (SBP-P5); GAP funding by National University of Singapore; AcRF Tier 2 by the Ministry of Education, Singapore; National Centre for Engineering Biology (NCEB), Singapore. We thank National University of Singapore HPC for providing GPU resources that facilitated the analysis. We thank Dr. Albert Xue Bo for providing valuable suggestions on ML modelling for enzyme engineering and Prof. Wen Shan Yew and Dr. Yan Pin Lim for sharing the Acella *E. coli* strain. We also extend our gratitude to Dr. Ruochong Zhang for insightful feedback during manuscript preparation.

## Author information

### Contributions

JY Zhang and CL Poh conceived the project and designed the PUSDA framework. JY Zhang implemented and benchmarked PUSDA, designed and conducted the experiments, generated data for ML, and analysed the results. CL Poh managed the project and provided the overall supervision. JY Zhang, JW Yeoh, and CL Poh jointly conceptualized the manuscript and developed its outline. JY Zhang led the writing of the manuscript, collected the results, and plotted the figures. Sangeetha processed the data for ML, trained ML models, and developed the feature selection approach. JW Yeoh retrieved the data from NGS and advised on ML analysis. JY Zhang and Sangeetha validated PUSDA. D Zheng, JR Goh, and ZY Lin assisted in conducting the sorting experiments and mutant characterization. All authors reviewed the manuscript.

## Ethics declarations

### Competing interests

A provisional patent related to the framework has been filed by National University of Singapore. The authors declare no other competing interests.

