## Supplementary information for "Machine learning-assisted enzyme engineering through ultra-high throughput sorting and large-scale sequence-function data generation"

### More information of FNS I

Flavone synthase I (FNS I) belongs to the family of flavone synthases, enzymes that catalyze the oxidation of flavanones to flavones, primarily found in plants. These enzymes were first discovered in 1975 by Sutter et al.[1], who observed that flavanone conversion to flavones required an oxygen-containing species and ferrous ion. Subsequent work by Britsch characterized FNS I as a soluble protein in parsley cells, with a molecular weight of approximately 48 kDa, an isoelectric point of pH 4.8, and a dependence on 2-oxoglutarate as the oxygen-containing species necessary for its reaction[2].

To date, other enzymes in this group, such as flavone synthase II, have been studied extensively, with some efforts directed at enhancing their activity and specificity through engineering[3,4]. However, FNS I remains relatively unexplored; existing studies are limited to basic characterizations, and no engineering attempts have been made to improve its function or elucidate its broader roles. As such, current knowledge of FNS I is restricted to its specific chemical reaction and phylogenetic or homologous relationships with other proteins.

The crystal structure of FNS I has not yet been resolved, and the AlphaFold-predicted structure provides only limited insights, with high confidence restricted to regions essential for enzyme functionality. Lower confidence is noted in regions with random coils, as well as at the C-terminus and N-terminus. Traditional enzyme engineering approaches typically rely on extensive prior knowledge, such as detailed 3D structure and active sites, which limits their applicability to less-characterized enzymes like FNS I.

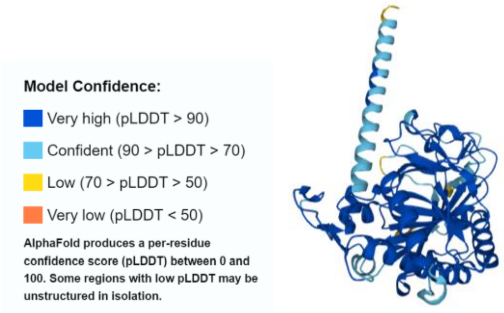

Figure S1: AlphaFold predicted model of FNS I enzyme with prediction confidence levels (from UniProt database) – the computationally predicted 3D structure is the only source of information about the structure of FNS I that can be found as there are no experimentally determined structures from FNS I.

**Biosensor dose response and characteristics**

To enable HTS for large mutant library, there is a need to find a high throughput detection method. There are a few detection methods that can be used to increase sorting throughput[5]. For example, electrochemical-Based Droplet Sorting was applied for NAD(P)-dependent oxidoreductase[6], however the throughput was limited to 30 per second. Mass Spectrometry has been coupled to HTS[7], but the throughput was limited to 33 per second. Raman-Activated Droplet Sorting was recently developed[8], however, the throughput was also limited, and the Raman signal can be easily interfered by the background noise. Directly detecting the target product will be more reliable in the assessment of enzyme performance, since focusing on co-factors, co-enzymes, or redox activity, like NAD/NADH, may overlook the entire reaction. In this work, we used a biosensor that directly detects apigenin concentration to conduct HTS and generate more reliable data for machine learning.

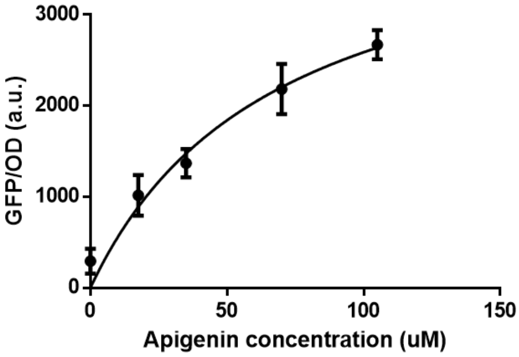

Figure S2: Characteristics of the apigenin biosensor. Apigenin biosensor showed an operational range from 17.5 to 105 μM, which covers the typical apigenin concentrations in the microbial production systems[9]. The $R^2$ reached 96.76%, indicating good fitting with Hill equation.

**Ultra-high throughput sorting platform**

In 2018, Nobel prize was awarded to Frances H. Arnold for inventing directed evolution, highlighting its significance and utility in enzyme engineering [10]. However, current screening methods heavily rely on labour effort, significantly limiting the throughput and efficiency. Synthetic biology offers promising approaches to implement the directed evolution for enzymes engineering [11]. The DBTL (Design-Build-Test-Learn) framework is commonly used in synthetic biology and metabolic engineering for developing biological systems. Despite its usefulness, DBTL approach faces challenges due to the complexity of biological systems and the lack of predictive design capabilities. At current practice, enzyme characterization and metabolite titration predominantly depend on HPLC or LC-MS methods, suffering from their labour-intensive operation and limited throughput.

High Throughput Sorting (HTS) is a powerful and automated technique to quickly test large numbers of samples [12,13]. Among different HTS techniques, FADS offer the advantages of miniaturization and compartmentalization, which are extremely suitable for ultra-HTS for enzymes or strains [14]. In this work, we setup customized FADS, as illustrated in Figure S3A-C, which consists of encapsulation, injection, and sorting modules. First, the single cell of mutant library was encapsulated into individual droplet to avoid the competition between different mutants, and encapsulation rate achieved more than 2000 droplets/s. Second, after 24 hours of cell growth, the substrate naringenin and inducer IPTG were injected into the droplet to induce the FNS enzyme expression and apigenin production, injection rate was more than 300 droplets/s. Third, after another 24 hours of apigenin production, the droplets were sorted based on the GFP intensity that emitted by the apigenin biosensor at a sorting rate of 200 droplets/s. Figure S1D showed the fluorescence images of droplets before and after sorting, in which the brighter droplets were sorted into positive channel, and the darker droplets were sorted into negative channel. To ensure the accuracy of the sorting, the GFP intensity of droplets in unsorted, negative, and positive groups were calculated. As seen in Figure S1E, the droplets in positive group indeed have high GFP intensity than these in negative groups, demonstrating high sorting accuracy (96%).

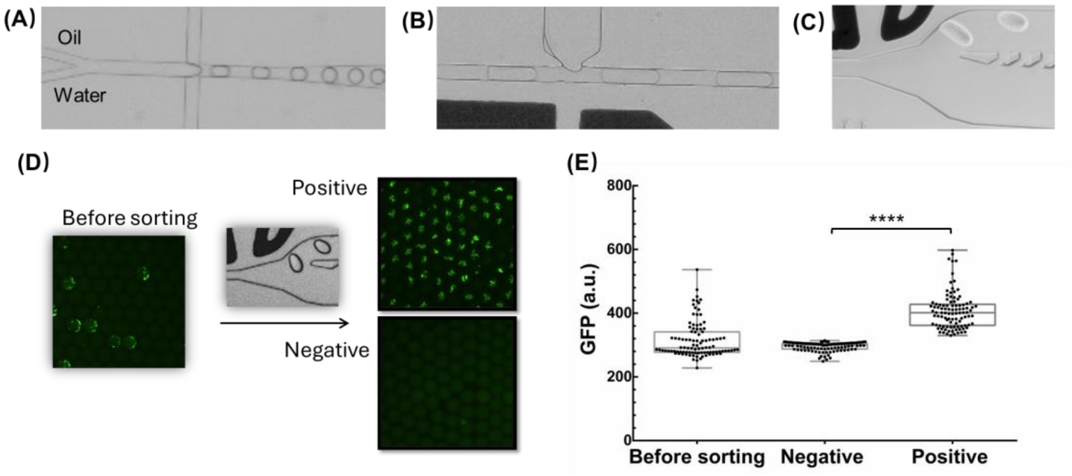

Figure S3: Ultra-high throughput sorting layout and performance. (A) Encapsulation. Single cell was encapsulated into individual droplet, at a rate of 2000/s. (B) Injection. After 24 hours incubation, the substrate and inducer were injected into the droplet, at a rate of 300/s. (C) Sorting. After another 24 hours incubation. The droplets were sorted into positive and negative channels, based on their GFP intensity. (D) The droplets before and after sorting. Almost all the droplets that sorted into positive had higher GFP intensity than negative groups. (E) The distribution of the sorted droplets' GFP intensity. Droplets with higher and lower GFP intensity were sorted into positive and negative groups. Data information: The experimental data were represented as mean ± S.D. (n = 100). Statistical significances of ****$P \leq 0.0001$, ***$P < 0.001$, **$P < 0.01$, and *$P < 0.05$ were calculated based on two-sample unpaired t-test.

### BLAST alignment

| | 0 | 1 | 2 | 3 | 4 | 5 | 6 | 7 | 8 | 9 | 10 | 11 | 12 | 13 | 14 | 15 | 16 | 17 | 18 | 19 | 20 | 21 | 22 | 23 | 24 | 25 | 26 | 27 | 28 | 29 | 30 | 31 | 32 |
|---|---|---|---|---|---|---|---|---|---|---|---|---|---|---|---|---|---|---|---|---|---|---|---|---|---|---|---|---|---|---|---|---|---|
| **0** | 0 | 0 | 0 | 0 | 0 | 0 | 0 | 0 | 0 | 0 | 0 | 0 | 0 | 0 | 0 | 0 | 0 | 0 | 0 | 0 | 0 | 0 | 0 | 0 | 0 | 0 | 0 | 0 | 0 | 0 | 0 | 0 | 0 |
| **1** | 0 | 0 | 0 | 0 | 0 | 0 | 0 | 0 | 0 | 0 | 0 | 0 | 0 | 0 | 0 | 0 | 0 | 0 | 0 | 0 | 0 | 0 | 0 | 0 | 0 | 0 | 0 | 0 | 0 | 0 | 0 | 0 | 0 |
| **2** | 0 | 0 | 0 | 0 | 0 | 0 | 0 | 0 | 0 | 0 | 0 | 0 | 0 | 0 | 0 | 0 | 0 | 0 | 0 | 0 | 0 | 0 | 0 | 0 | 0 | 0 | 0 | 0 | 0 | 0 | 0 | 0 | 0 |
| **3** | 0 | 0 | 0 | 0 | 0 | 0 | 0 | 0 | 0 | 0 | 0 | 0 | 0 | 0 | 0 | 0 | 0 | 0 | 0 | 0 | 0 | 0 | 0 | 0 | 0 | 0 | 0 | 0 | 0 | 0 | 0 | 0 | 0 |
| **4** | 0 | 0 | 0 | 0 | 0 | 0 | 0 | 0 | 0 | 0 | 0 | 0 | 0 | 0 | 0 | 0 | 0 | 0 | 0 | 0 | 0 | 0 | 0 | 0 | 0 | 0 | 0 | 0 | 0 | 0 | 0 | 4 | 0 |
| **5** | 0 | 0 | 0 | 0 | 0 | 0 | 0 | 0 | 0 | 0 | 0 | 0 | 0 | 0 | 0 | 0 | 0 | 0 | 0 | 0 | 0 | 0 | 0 | 0 | 0 | 0 | 0 | 0 | 0 | 0 | 0 | 0 | 0 |
| **6** | 0 | 0 | 0 | 0 | 0 | 0 | 0 | 0 | 0 | 0 | 0 | 0 | 0 | 0 | 0 | 0 | 0 | 0 | 0 | 0 | 0 | 0 | 0 | 0 | 0 | 0 | 0 | 0 | 0 | 0 | 0 | 0 | 0 |
| **7** | 0 | 0 | 0 | 0 | 0 | 0 | 0 | 0 | 0 | 0 | 0 | 0 | 0 | 0 | 0 | 0 | 0 | 0 | 0 | 0 | 0 | 0 | 0 | 0 | 0 | 0 | 0 | 0 | 0 | 0 | 0 | 0 | 0 |
| **8** | 0 | 0 | 0 | 0 | 0 | 0 | 0 | 0 | 0 | 0 | 0 | 0 | 0 | 0 | 0 | 0 | 0 | 0 | 0 | 0 | 0 | 0 | 0 | 0 | 0 | 0 | 0 | 0 | 0 | 0 | 0 | 0 | 0 |
| **9** | 0 | 0 | 0 | 0 | 0 | 0 | 0 | 0 | 0 | 0 | 0 | 0 | 0 | 0 | 0 | 0 | 0 | 0 | 0 | 0 | 0 | 0 | 0 | 0 | 0 | 0 | 0 | 0 | 0 | 0 | 0 | 0 | 0 |
| **10** | 0 | 0 | 0 | 0 | 0 | 0 | 0 | 0 | 0 | 0 | 0 | 0 | 0 | 0 | 0 | 0 | 0 | 0 | 0 | 0 | 0 | 0 | 0 | 0 | 0 | 0 | 0 | 0 | 0 | 0 | 0 | 0 | 0 |
| **11** | 0 | 0 | 0 | 0 | 0 | 0 | 0 | 0 | 0 | 0 | 0 | 0 | 0 | 0 | 0 | 0 | 0 | 0 | 0 | 0 | 0 | 0 | 0 | 0 | 0 | 0 | 0 | 0 | 0 | 0 | 0 | 0 | 0 |
| **12** | 0 | 0 | 0 | 0 | 0 | 0 | 0 | 0 | 0 | 0 | 0 | 0 | 0 | 0 | 0 | 0 | 0 | 0 | 0 | 0 | 0 | 0 | 0 | 0 | 0 | 0 | 0 | 0 | 0 | 0 | 0 | 0 | 0 |
| **13** | 0 | 0 | 0 | 0 | 0 | 0 | 0 | 0 | 0 | 0 | 0 | 0 | 0 | 0 | 0 | 0 | 0 | 0 | 0 | 0 | 0 | 0 | 0 | 0 | 0 | 0 | 0 | 0 | 0 | 0 | 0 | 0 | 0 |
| **14** | 0 | 0 | 0 | 0 | 0 | 0 | 0 | 0 | 0 | 0 | 0 | 0 | 0 | 0 | 0 | 0 | 0 | 0 | 0 | 0 | 0 | 0 | 0 | 0 | 0 | 0 | 0 | 0 | 0 | 0 | 0 | 0 | 0 |
| **15** | 0 | 0 | 0 | 0 | 0 | 0 | 0 | 0 | 0 | 0 | 0 | 0 | 0 | 0 | 0 | 0 | 0 | 0 | 0 | 0 | 0 | 0 | 0 | 0 | 0 | 0 | 0 | 0 | 0 | 0 | 0 | 0 | 0 |
| **16** | 0 | 0 | 0 | 0 | 0 | 0 | 0 | 0 | 0 | 0 | 0 | 0 | 0 | 0 | 0 | 0 | 0 | 0 | 0 | 0 | 0 | 0 | 0 | 0 | 0 | 0 | 0 | 0 | 0 | 0 | 0 | 0 | 0 |
| **17** | 0 | 0 | 0 | 0 | 0 | 0 | 0 | 0 | 0 | 0 | 0 | 0 | 0 | 0 | 0 | 0 | 0 | 0 | 0 | 0 | 0 | 0 | 0 | 0 | 0 | 0 | 0 | 0 | 0 | 0 | 0 | 0 | 0 |
| **18** | 0 | 0 | 0 | 0 | 0 | 0 | 0 | 0 | 0 | 0 | 0 | 0 | 0 | 0 | 0 | 0 | 0 | 0 | 0 | 0 | 0 | 0 | 0 | 0 | 0 | 2 | 0 | 0 | 0 | 0 | 0 | 0 | 0 |
| **19** | 0 | 0 | 0 | 0 | 0 | 0 | 0 | 0 | 0 | 0 | 0 | 0 | 0 | 0 | 0 | 0 | 0 | 0 | 0 | 0 | 0 | 0 | 0 | 0 | 0 | 0 | 0 | 0 | 0 | 0 | 0 | 0 | 0 |
| **20** | 0 | 0 | 0 | 0 | 0 | 0 | 0 | 0 | 0 | 0 | 0 | 0 | 0 | 0 | 0 | 0 | 0 | 0 | 0 | 0 | 0 | 0 | 0 | 0 | 0 | 0 | 0 | 0 | 0 | 0 | 0 | 0 | 0 |
| **%Sub to A** | 0.0 | 0.0 | 0.0 | 0.0 | 0.0 | 0.0 | 0.0 | 0.0 | 0.0 | 0.0 | 0.0 | 0.0 | 0.0 | 0.0 | 0.0 | 0.0 | 0.0 | 0.0 | 0.0 | 0.0 | 0.0 | 0.0 | 0.0 | 0.0 | 0.0 | 0.0 | 0.0 | 0.0 | 0.0 | 0.0 | 0.0 | 0.0 | 0.0 |
| **%Sub to T** | 0.0 | 0.0 | 0.0 | 0.0 | 0.0 | 0.0 | 0.0 | 0.0 | 0.0 | 0.0 | 0.0 | 0.0 | 0.0 | 0.0 | 0.0 | 0.0 | 0.0 | 0.0 | 0.0 | 0.0 | 0.0 | 0.0 | 0.0 | 0.0 | 0.0 | 4.545455 | 0.0 | 0.0 | 0.0 | 0.0 | 0.0 | 0.0 | 0.0 |
| **%Sub to C** | 0.0 | 0.0 | 0.0 | 0.0 | 0.0 | 0.0 | 0.0 | 0.0 | 0.0 | 0.0 | 0.0 | 0.0 | 0.0 | 0.0 | 0.0 | 0.0 | 0.0 | 0.0 | 0.0 | 0.0 | 0.0 | 0.0 | 0.0 | 0.0 | 0.0 | 0.0 | 0.0 | 0.0 | 0.0 | 0.0 | 0.0 | 0.0 | 0.0 |
| **%Sub to G** | 0.0 | 0.0 | 0.0 | 0.0 | 0.0 | 0.0 | 0.0 | 0.0 | 0.0 | 0.0 | 0.0 | 0.0 | 0.0 | 0.0 | 0.0 | 0.0 | 0.0 | 0.0 | 0.0 | 0.0 | 0.0 | 0.0 | 0.0 | 0.0 | 0.0 | 0.0 | 0.0 | 0.0 | 0.0 | 0.0 | 0.0 | 4.166667 | 0.0 |
| **%Insertion** | 0.0 | 0.0 | 0.0 | 0.0 | 0.0 | 0.0 | 0.0 | 0.0 | 0.0 | 0.0 | 0.0 | 0.0 | 0.0 | 0.0 | 0.0 | 0.0 | 0.0 | 0.0 | 0.0 | 0.0 | 0.0 | 0.0 | 0.0 | 0.0 | 0.0 | 0.0 | 0.0 | 0.0 | 0.0 | 0.0 | 0.0 | 0.0 | 0.0 |
| **%Deletion** | 0.0 | 0.0 | 0.0 | 0.0 | 0.0 | 0.0 | 0.0 | 0.0 | 0.0 | 0.0 | 0.0 | 0.0 | 0.0 | 0.0 | 0.0 | 0.0 | 0.0 | 0.0 | 0.0 | 0.0 | 0.0 | 0.0 | 0.0 | 0.0 | 0.0 | 0.0 | 0.0 | 0.0 | 0.0 | 0.0 | 0.0 | 0.0 | 0.0 |

Figure S4: BLAST alignment. Parsing of alignment to get specific substitution types across samples. BLAST has the advantages to handle large dataset and find homologous sequences in databases. Therefore, sequence alignment was done using BLAST to obtain information such as the sequence length, region of match, number of gaps, positions of mismatches, and mutation events.

### Data filtration/Training data curation

Mutation events that result in truncated enzymes were discarded, as they could introduce noise and potentially bias downstream analysis. This allows filtering of the samples to only include full length matches and only substitution events (no gaps), excluding the sequencing error, insertion and deletion, and truncation events, which might result in low model accuracy.

It should be noted that although only substitution events for full length matched samples were being considered, there could be cases of insertions or deletions that did not result in a length changing mutation (especially when the events of insertions and deletions happen simultaneously to compensate for each other). On the other hand, some substitutions could introduce stop codons, resulting in truncated enzymes. However, this study would like to first focus on substitution events only so that important mutations that are linked to the function changes can be identified. As a result, insertions and deletions were being ignored at this stage. To incorporate this level of scrutiny to the filtering of dataset, additional computational screening steps were required to ensure that the insertions and deletions were not changing the overall length of the sequence. As we were dealing with a million samples, this could be time and computation power-consuming. Including sequences that can lead to truncated enzymes was not beneficial to this study as these samples could be noise and could skew downstream analysis. Thus, only substitution events were being considered.

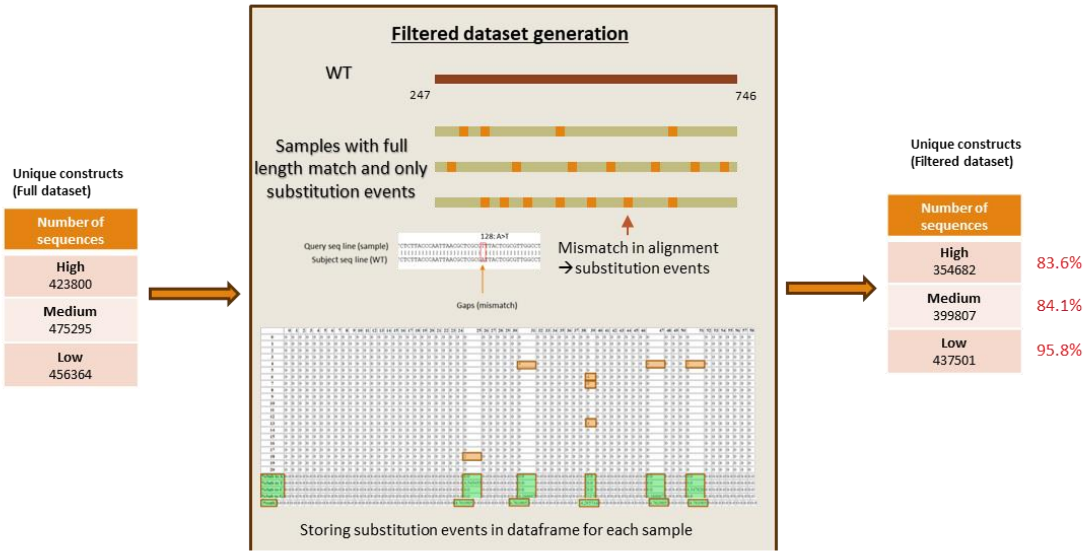

Figure S1: Data filtration. BLAST alignment was used to filter out faulty reads, overlapping sequences, and truncated sequences. Initially, faulty reads were discarded, followed by the removal of duplicate sequences to retain only unique ones. Truncated sequences were subsequently eliminated, resulting in final sequence counts of 354,682 for the High group, 399,807 for the Medium group, and 437,501 for the Low group.

### DNA landscape for H, M, L groups

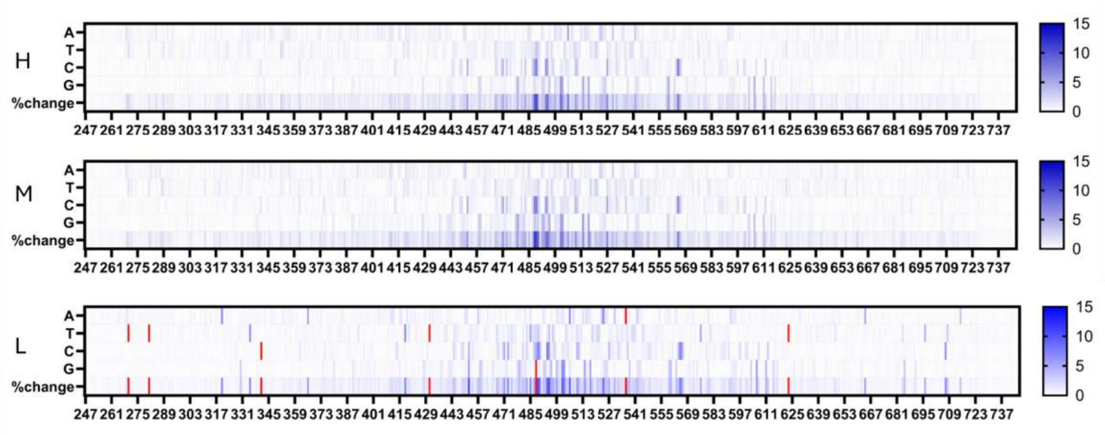

Figure S6: DNA landscape of the sorted mutant library. To visualize the DNA composition of the sorted mutant library, the DNA landscapes of the mutants in the H, M, and L groups are plotted.

### Occurrence of DNA mutation events

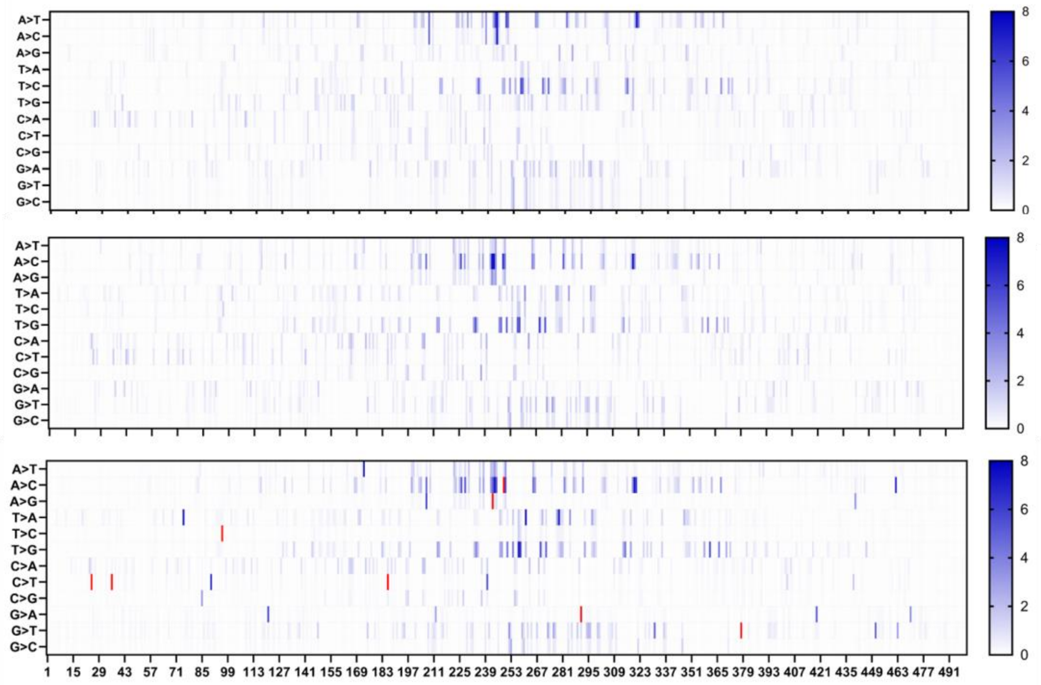

Figure S7: DNA mutation events landscape of the sorted mutant library. H, M, L. To visualize the distribution of specific mutation events, the landscapes of specific mutation events were also plotted. Overall, the mutation events in L had higher occurrence than H and M. A to T was the most occurring in H, and most of the mutations happened at location of 241, 252, and 320. A to C was most occurring in M, through the positions of 241, 250, and 313. C to T was the most occurring mutation event in L, at positions of 22, 35, 88, 185, and 240.

### Specific AA mutation events in H, M, L groups

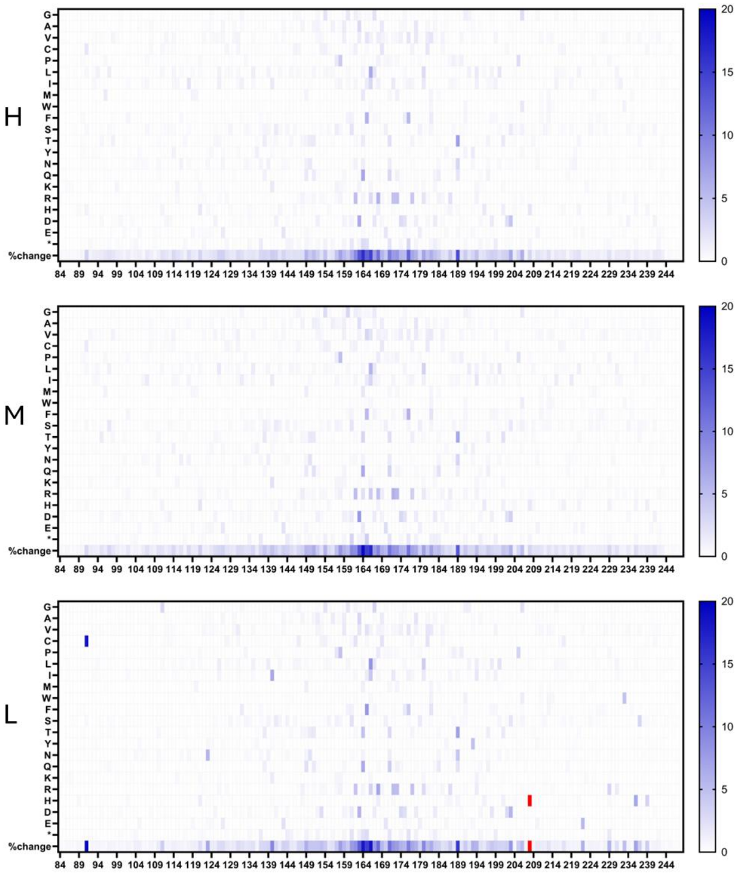

Figure S8: Specific AA mutation events. From the AA mutation events landscapes, we observed that high occurring mutation events (>5% occurrence) happened in H were 163D, 164Q, 165F, 166L, 168R, 176F, and 189T. The most abundant mutations event happened in M were 162K, 163D, 164Q, 166L, 172R, 173R, 177F, and 189T. The most abundant mutations event happened in L were 89C, 123N, 140I, 163D, 164Q, 165F, 166L, 168R, 172R, 173R, 189T, 203D, 208H, 222E, and 236H.

### Mutation polarity distribution

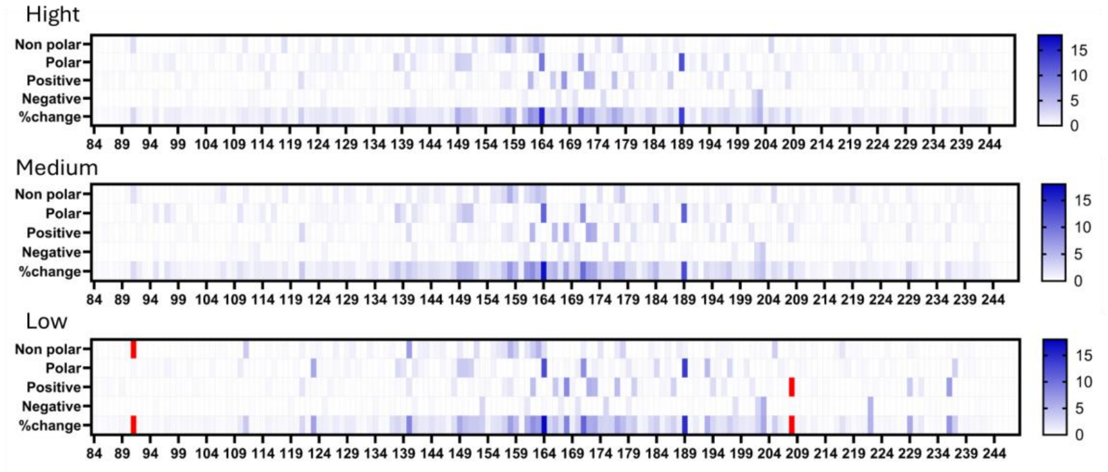

Figure S9: Amino acid polarity and charge distribution of sorted mutant library. H and M had similar polarity and charge composition, and AA with more property change located at AA164 and AA189. While the AA that with more property change located at AA91 and AA208.

### ML models training

We trained several ML models, including Logistic Regression, Random Forest, and Support Vector Machine on exploratory data (30k samples) and used an ensemble voting classifier, based on majority rule, to gain insights into model performance and optimize hyperparameters. All models achieved high accuracy (~0.85) on the training set (Figure S10A). Evaluation metrics, including a confusion matrix on validation data, indicated generally accurate classification, with some misclassification between classes M and L, likely due to their similar mutation patterns (Figures S10B-C). Key mutations and positions for experimental follow-up were identified by ranking important features using the classifier (Figure S10D). Additionally, a Multilayer Perceptron (MLP) was trained on the full dataset (over 1 million samples), achieving high accuracy (>0.85) on both training and validation sets without significant overfitting (Figure S10F).

Initially, we trained models on the full dataset without filtering to assess their effectiveness on a one-hot encoded DNA sequence input. Subsequent modifications, like filtering based on BLAST alignment or using one-hot encoded amino acid sequences, were not expected to drastically alter model performance. This phase aimed to clarify why certain models outperformed others on this dataset, especially since no standard model exists in the literature for this data representation.

Notably, the Multilayer Perceptron performed comparably to Random Forest. However, the MLP requires more intensive tuning of intermediate layers, which can be resource-demanding, particularly with large datasets. Random Forest was favoured for its simpler structure and ease of interpretation compared to Neural Networks, which require significant adjustments to intermediate layers and lack built-in support for such tuning in scikit-learn. Moreover, Random Forest allows straightforward feature importance ranking, while MLP requires additional calculations to interpret feature impacts.

Random Forest, pioneered by Leo Breiman and Adele Cutler, aggregates predictions from multiple decision trees with built-in randomness, reducing correlation among trees. The one-hot encoded format enables clear binary splits at each node, guiding tree growth primarily in a single direction. This straightforward structure may contribute to the model's robust performance.

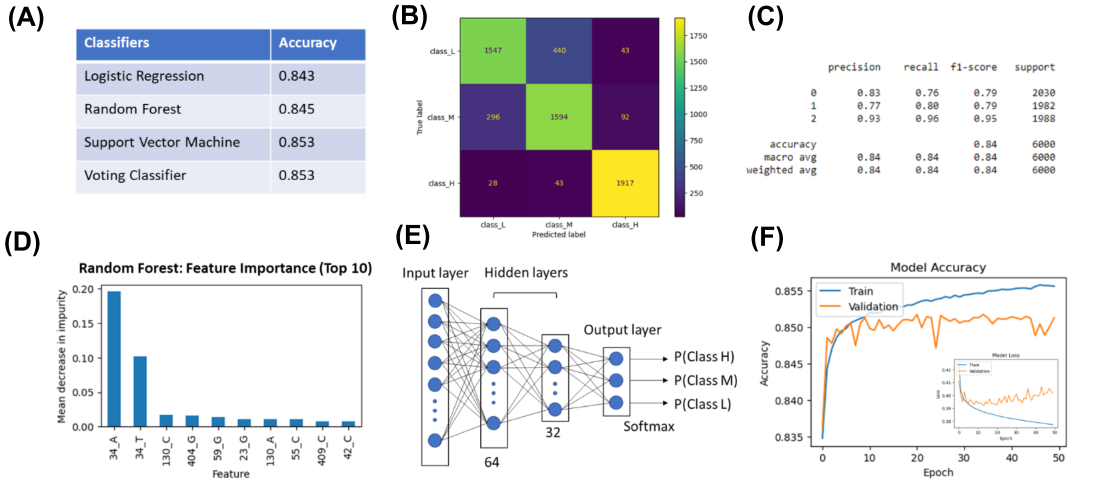

Figure S10**:** ML model training. (A) Model performance for the three models and a voting classifier based on the three classifiers. (B) Confusion matrix analysis. (C) Classification report for the three-evaluation metrics (precision, recall, and f1-score). (D) The top 10 important features identified using the Random Forest model. (E) Multilayer Perceptron model architecture. (F) Model accuracy and loss during the training and validation phases.

### One-hot encoding

One-hot encoding is chosen as the DNA sequence representation method based on the presence or absence of a particular base at each position. This serves as a suitable depiction of the mutant samples, enabling additional analysis to ascertain the importance of each base at a specific position in enhancing the model's predictive accuracy. This will also minimize the input rather than representing each base a series of matrix. On the other hand, the output for the models is categorical, ranking the three functional groups. This introduces a ranking among the output variables, which differs from the input variable that only capture the presence or absence of sequence at specific positions.

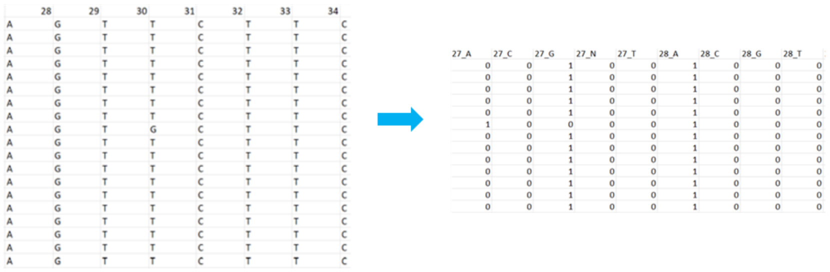

Figure S11: One-hot encoding. The unique sequences of the mutants were then subjected to one-hot encoding, converting nucleotide or amino acid sequences into binary matrices for downstream computational analysis.

### Model evaluation

Overall, the models' precision, recall, F1-score, and accuracy except for GNB were all higher than 0.8, while for M and H groups were lower but most of the scores were higher than 0.6. For the precision, recall, and F1-score, other than SGD, GTB, AC, and NGB, the rest models achieved scores higher than 0.6, among which RFC performed best for all the H, M, and L groups. As for accuracy, RFC, EC, and MLP performed the best among all the models, achieving accuracy higher than 80%. The other models had varying performances ranging from as low as 38% (GNB) to 77% (HBGB) accuracy on the test dataset. GNB performed the worse, likely due to its feature independence and normal distribution assumption[15].

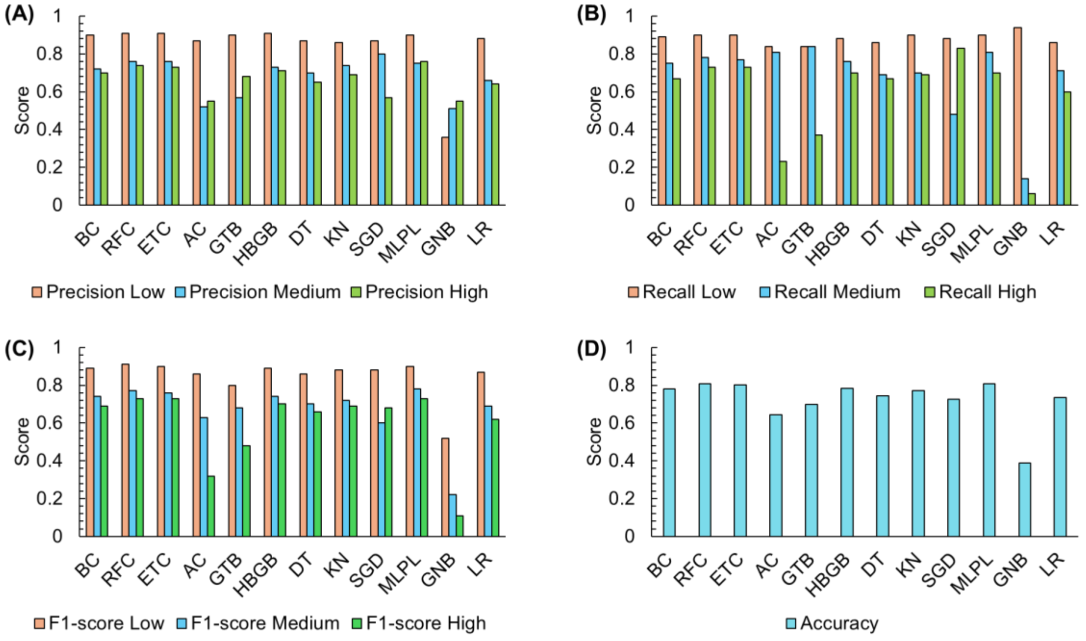

Figure S12: comparison of different ML models. (A) Precision score of different models. RFC, ETC, and MLPL were higher than the rest. (B) Recall score of different models. BC, RFC, and ETC had higher recall scores than the rest. (C) F1-score of different models. BC, RFC, ETC, and MLPL were higher than the rest. (D) Accuracy of different models. RFC, ETC, and MLPL were higher than the rest. Overall, RFC models performed best among these models.

**Evaluating DNA and AA sequence as input for ML**

The accuracy of the model using DNA sequence as input was 0.81, whereas it was 0.72 for AA input when comparing all the H, M, and L groups. Similarly, when comparing between only two groups (H-L, H-M, M-L) separately, the precision, recall, and F1-scores were also higher when using DNA sequences as input. The highest model accuracy reached 0.932 and 0.935 when comparing H-L and M-L, respectively. However, the accuracy was only 0.77 when comparing H-M. This indicates that the model was better at distinguishing L from H and M groups.

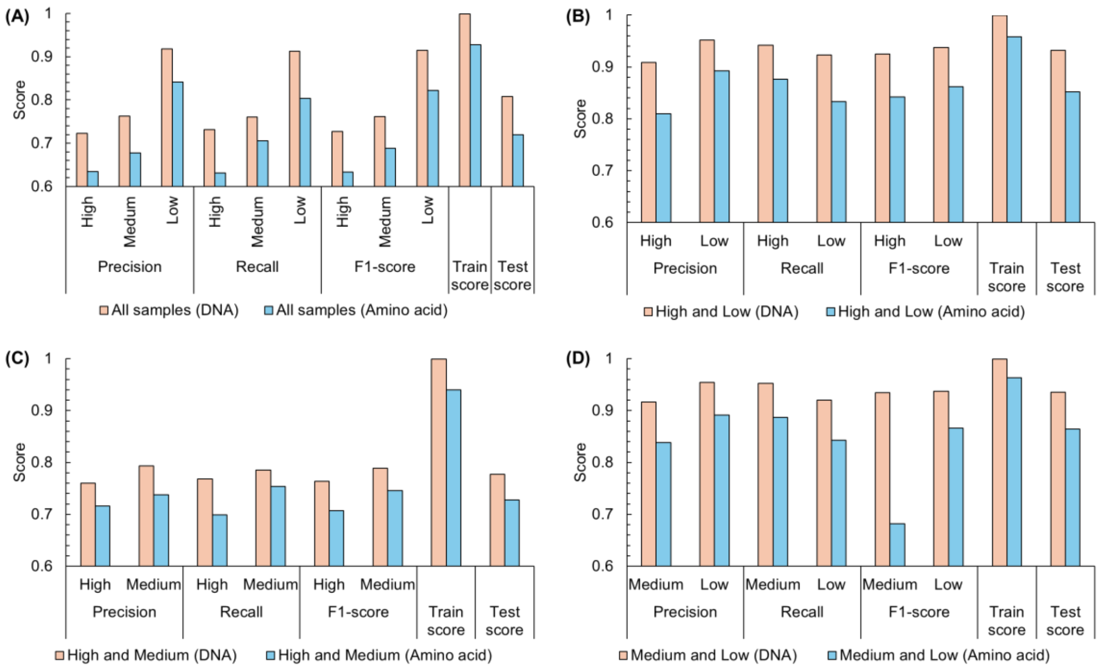

Figure S13: comparison of RFC model performance when using DNA or AA as input. (A) RFC performance when including H, M, and L as input. Using DNA as input outperformed AA. (B) RFC performance when only including H and L as input. Using DNA as input outperformed AA. (C) RFC performance when only including H and M as input. Using DNA as input outperformed AA. (D) RFC performance when only including M and L as input. Using DNA as input outperformed AA. Overall, using DNA as input achieved better model performance throughout the four conditions. Besides, the model performance maximized when only including H-L or M-L as input.

### Feature importance

The RFC model was used to extract the features, the runs that included Low in the input (High & Low, Medium & Low) generated higher feature importance and thus they were better in distinguishing the two groups. However, when only High & Medium were included as the input, the feature importance had lower values, suggesting that it is more difficult to identify important features to distinguish between the High and Medium groups. This reinforced to couple the feature importance with findings from the mutational landscape observed in BLAST analysis. Thus, there is a need to delve into alternative representation of mutation to discern which mutations are most important in affecting the performance of each group.

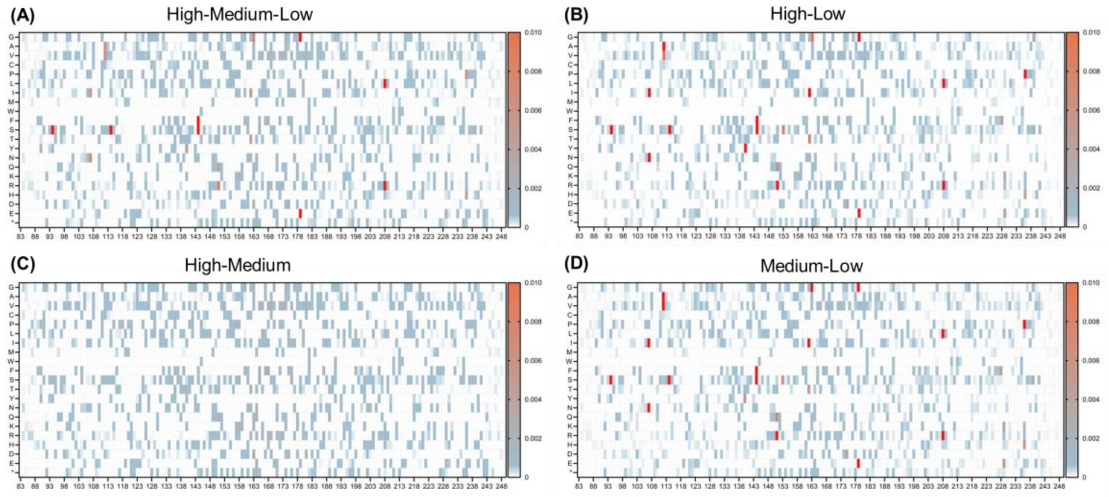

Figure S14: Feature importance landscape. (A) Landscape of feature importance when using H, M, and L groups as input. (B) Landscape of feature importance when using H and L groups as input. (C) Landscape of feature importance when using H and M groups as input. (D) Landscape of feature importance when using M and L groups as input.

### Distribution of the feature importances

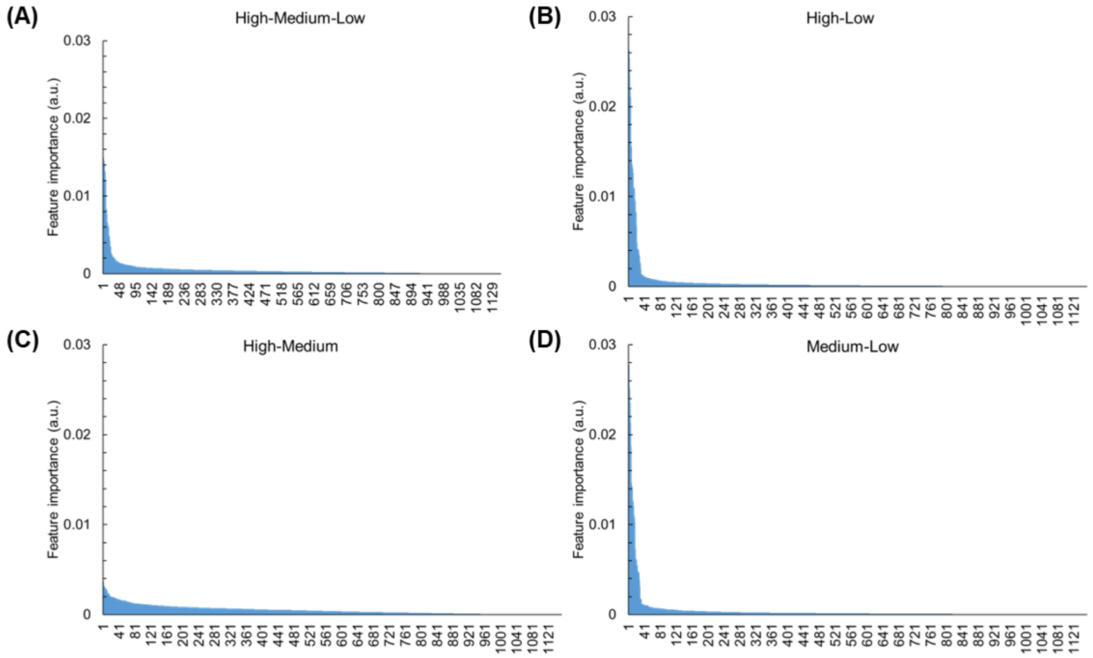

Figure S15: Distribution of feature importance. (A) Distribution of feature importance when using H, M, and L groups as input. (B) Distribution of feature importance when using H and L groups as input. (C) Distribution of feature importance when using H and M groups as input. The overall feature importance is much less as compared to other input conditions. (D) Distribution of feature importance when using M and L groups as input. Only part of the features (top 30%) has relatively higher importance score (>0.0002). The x-axis is the number of the features generated by the ML model.

### Top 30% feature importances

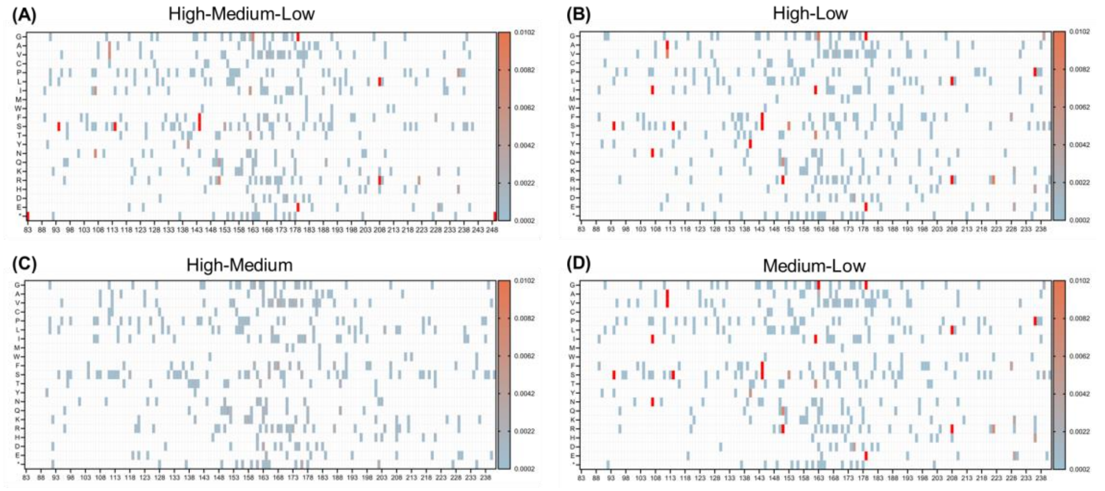

Figure S16: Landscape of top 30% feature importance. (A) Landscape of top 30% feature importance when using H, M, and L groups as input. (B) Landscape of top 30% feature importance when using H and L groups as input. (C) Landscape of top 30% feature importance when using H and M groups as input. (D) Landscape of top 30% feature importance when using M and L groups as input.

### BLAST mutation events (more than 1% occurrence)

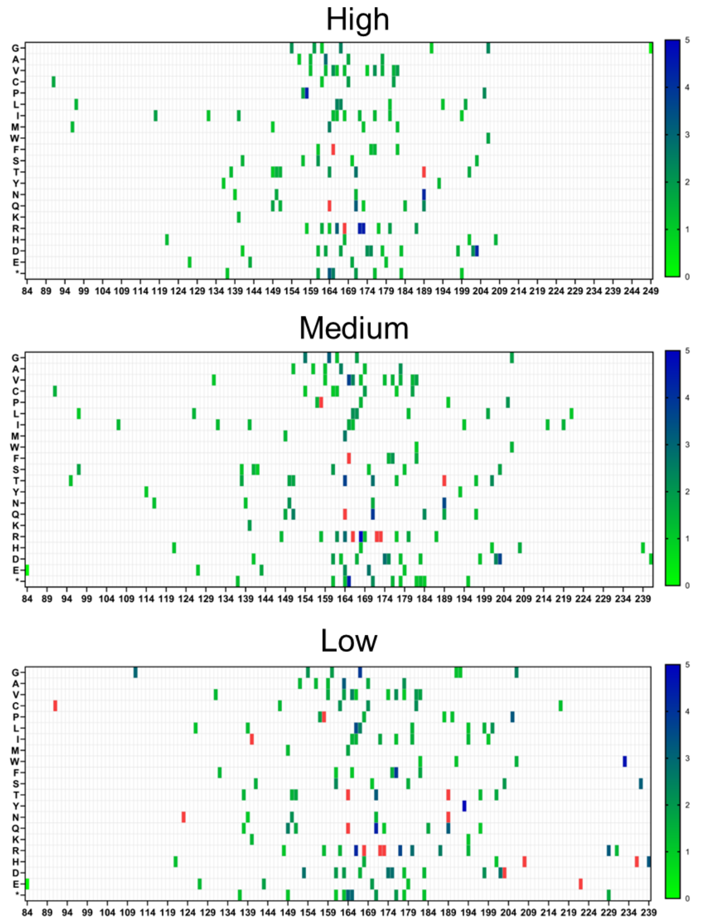

Figure S17: Landscape of mutation events that had more than 1% occurrence. L group has overall higher mutation events occurrence spanning over the whole search space, while H and M groups' mutation events clustered at the middle region.

### Selected 504 features

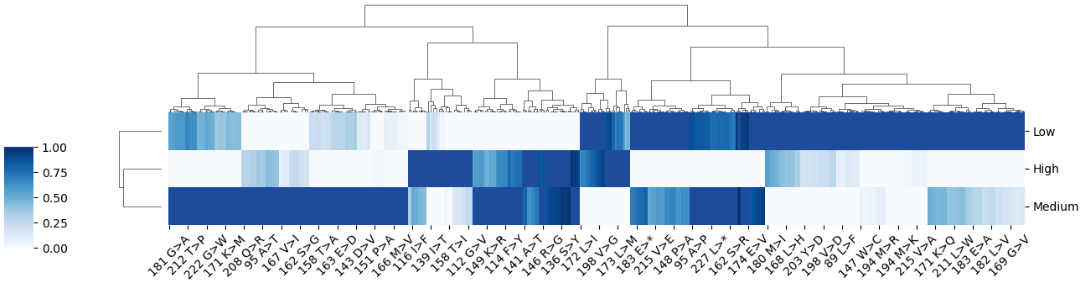

Figure S18: Clustered heatmap of selected 504 features obtained by intersecting ML-generated features and high-occurrence mutation events. Mutation occurrences were normalized to a scale from 0 to 1 within each performance group. The clustering analysis revealed significant mutation occurrences across groups, thus enhancing the distinction between the three performance groups. This visualization highlights the general trends in mutation occurrences across performance groups, but it cannot identify most critical mutations contribute to the characteristics of each group. Individual mutations may exhibit low absolute frequencies across all groups, making it difficult to identify crucial mutations from this heatmap alone. Furthermore, while the heatmap provides valuable insights into single mutation occurrences, it does not capture the potential relevance of joint mutation occurrences, underscoring the need to analyse the mutations in combination.

### Mutation combinations

To explore more complex higher-dimensional mutation events, we also studied the mutation events in combination. The mutant samples, which had previously been represented by DNA or AA sequences, were redefined based on the presence or absence of the selected 504 mutation events, generating a presence/absence matrix of mutations for each sample across all three performance groups.

(A)

504 mutation events

| Sample | 89 L>F | 89 L>I | ... | 241 G>S | 241 G>R |
|---|---|---|---|---|---|
| H_1 | 0 | 1 | | 0 | 1 |
| H_2 | 0 | 0 | | 1 | 0 |
| . . . | | | | | |
| H_354681 | 1 | 0 | | 0 | 0 |
| H_354682 | 0 | 0 | | 0 | 1 |

(B)

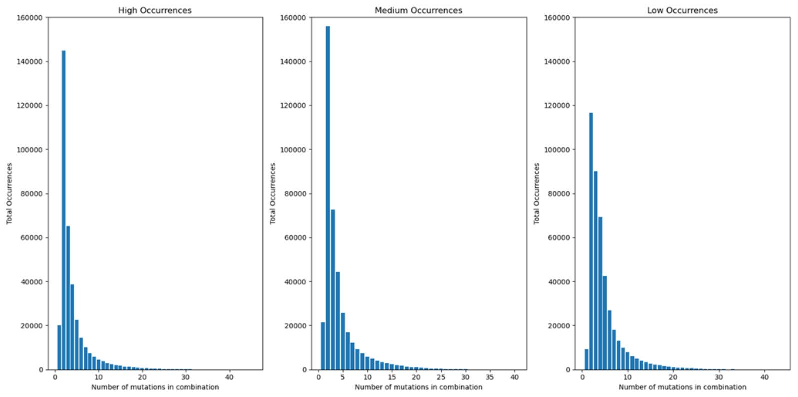

(C)

High

| Mutation list | Samples | Percentage |
|---|---|---|
| () | 20102 | 5.667612 |
| ('164 K>Q',) | 1259 | 0.354966 |
| ('189 K>T',) | 1250 | 0.352428 |
| ('166 M>L',) | 1059 | 0.298577 |
| ('163 E>D',) | 975 | 0.274894 |
| ('165 L>F',) | 975 | 0.274894 |
| ('176 L>F',) | 921 | 0.259669 |

Medium

| Mutation list | Samples | Percentage |
|---|---|---|
| () | 21428 | 5.359586 |
| ('163 E>D',) | 1374 | 0.343666 |
| ('164 K>Q',) | 1374 | 0.343666 |
| ('166 M>L',) | 1079 | 0.26988 |
| ('189 K>T',) | 1068 | 0.267129 |
| ('173 L>R',) | 940 | 0.235113 |
| ('165 L>F',) | 938 | 0.234613 |

Low

| Mutation list | Samples | Percentage |
|---|---|---|
| () | 9197 | 2.102167 |
| ('208 Q>H',) | 8194 | 1.87291 |
| ('91 R>C',) | 3418 | 0.781255 |
| ('222 G>E', '236 Q>H') | 1675 | 0.382856 |
| ('91 R>C', '208 Q>H') | 1595 | 0.364571 |
| ('123 D>N', '140 N>I') | 1410 | 0.322285 |
| ('193 D>Y', '233 G>W') | 1371 | 0.313371 |
| ('153 G>D', '237 A>S') | 937 | 0.214171 |

Figure S19: Dissecting higher-dimensional features. (A) Workflow for creating mutation combinations. (B) Distribution of mutation combinations based on the number of single mutation events. (C) Occurrences of mutation combinations in each performance group.

### Ratio-based selection approach

| Feature importance | Features | DNA position | WT base | Changed base | DNA substituion | Feature_DNA | AA position | WT codon | WT AA | Changed codon | Changed AA | AA Substitution | Feature_AA | |
|---|---|---|---|---|---|---|---|---|---|---|---|---|---|---|
| 0.07267773 | 24_T | 270 | C | T | C>T | 270 C>T | 90 | TCC | S | TCT | S | S>S | 90 S>S | |
| 0.055308802 | 24_C | 270 | C | C | C>C | 270 C>C | 90 | TCC | S | TCC | S | S>S | 90 S>S | |
| 0.030917288 | 95_C | 341 | T | C | T>C | 341 T>C | 114 | TTT | F | TCT | S | F>S | 114 F>S | 1 |
| 0.028294156 | 35_T | 281 | T | T | T>T | 281 T>T | 94 | TTC | F | TTC | F | F>F | 94 F>F | |
| 0.027758493 | 290_A | 536 | C | A | C>A | 536 C>A | 179 | GCG | A | GAG | E | A>E | 179 A>E | 2 |
| 0.025266345 | 377_T | 623 | A | T | A>T | 623 A>T | 208 | CAG | Q | CTG | L | Q>L | 208 Q>L | 3 |
| 0.024995702 | 290_G | 536 | C | G | C>G | 536 C>G | 179 | GCG | A | GGG | G | A>G | 179 A>G | 4 |
| 0.023524678 | 185_T | 431 | A | T | A>T | 431 A>T | 144 | TAC | Y | TTC | F | Y>F | 144 Y>F | 5 |
| 0.02137588 | 35_C | 281 | T | C | T>C | 281 T>C | 94 | TTC | F | TCC | S | F>S | 94 F>S | 6 |
| 0.018735157 | 377_G | 623 | A | G | A>G | 623 A>G | 208 | CAG | Q | CGG | R | Q>R | 208 Q>R | 7 |
| 0.018200717 | 95_T | 341 | T | T | T>T | 341 T>T | 114 | TTT | F | TTT | F | F>F | 114 F>F | |
| 0.015050933 | 120_G | 366 | A | G | A>G | 366 A>G | 122 | GGA | G | GGG | G | G>G | 122 G>G | |
| 0.014675406 | 185_C | 431 | A | C | A>C | 431 A>C | 144 | TAC | Y | TCC | S | Y>S | 144 Y>S | 8 |
| 0.014317091 | 74_T | 320 | C | T | C>T | 320 C>T | 107 | ACT | T | ATT | I | T>I | 107 T>I | 9 |
| 0.014145186 | 206_G | 452 | C | G | C>G | 452 C>G | 151 | CCA | P | CGA | R | P>R | 151 P>R | 10 |
| 0.012726046 | 89_T | 335 | G | T | G>T | 335 G>T | 112 | GGC | G | GTC | V | G>V | 112 G>V | 11 |
| 0.012523011 | 461_C | 707 | A | C | A>C | 707 A>C | 236 | CAA | Q | CCA | P | Q>P | 236 Q>P | 12 |
| 0.012509939 | 120_A | 366 | A | A | A>A | 366 A>A | 122 | GGA | G | GGA | G | G>G | 122 G>G | |
| 0.01208744 | 89_C | 335 | G | C | G>C | 335 G>C | 112 | GGC | G | GCC | A | G>A | 112 G>A | 13 |
| 0.011030609 | 239_T | 485 | G | T | G>T | 485 G>T | 162 | AGC | S | ATC | I | S>I | 162 S>I | 14 |
| 0.010802877 | 242_G | 488 | A | G | A>G | 488 A>G | 163 | GAA | E | GGA | G | E>G | 163 E>G | 15 |
| 0.010258324 | 74_A | 320 | C | A | C>A | 320 C>A | 107 | ACT | T | AAT | N | T>N | 107 T>N | 16 |
| 0.009889449 | 461_A | 707 | A | A | A>A | 707 A>A | 236 | CAA | Q | CAA | Q | Q>Q | 236 Q>Q | |
| 0.009507811 | 418_G | 664 | G | G | G>G | 664 G>G | 222 | GGG | G | GGG | G | G>G | 222 G>G | |
| 0.00803435 | 206_A | 452 | C | A | C>A | 452 C>A | 151 | CCA | P | CAA | Q | P>Q | 151 P>Q | 17 |
| 0.008005149 | 84_G | 330 | A | G | A>G | 330 A>G | 110 | AAA | K | AAG | K | K>K | 110 K>K | |
| 0.007447649 | 172_A | 418 | A | A | A>A | 418 A>A | 140 | AAC | N | AAC | N | N>N | 140 N>N | |
| 0.007258934 | 418_A | 664 | G | A | G>A | 664 G>A | 222 | GGG | G | AGG | R | G>R | 222 G>R | 18 |
| 0.006942263 | 450_T | 696 | T | T | T>T | 696 T>T | 232 | GTT | V | GTT | V | V>V | 232 V>V | |
| 0.006775495 | 242_A | 488 | A | A | A>A | 488 A>A | 163 | GAA | E | GAA | E | E>E | 163 E>E | |
| 0.006660629 | 330_T | 576 | A | T | A>T | 576 A>T | 192 | GTA | V | GTT | V | V>V | 192 V>V | |
| 0.00624808 | 211_A | 457 | G | A | G>A | 457 G>A | 153 | GGT | G | AGT | S | G>S | 153 G>S | 19 |
| 0.006165863 | 239_C | 485 | G | C | G>C | 485 G>C | 162 | AGC | S | ACC | T | S>T | 162 S>T | 20 |
| 0.006019531 | 172_T | 418 | A | T | A>T | 418 A>T | 140 | AAC | N | TAC | Y | N>Y | 140 N>Y | |
| 0.005968765 | 450_G | 696 | T | G | T>G | 696 T>G | 232 | GTT | V | GTG | V | V>V | 232 V>V | |

Figure S20: Ratio-based higher dimensional feature selection approach. The ratio of these mutation combinations between each group in each category will then be calculated to identify those with a higher prevalence in each performance group. Also, unique mutation combinations that are solely present in each group are also identified.

### Analysis of local effects of amino acid changes

The 3D structural image illustrates the positions and types of mutations from the generated list. Mutations occur at various locations across the protein, with many positioned on the surface where they interact with the external environment. Notably, mutations such as L96M and P97L are clustered in proximity in the High configuration, while R91C appears near this region but in the Low configuration. Similarly, G181V, E183K, and K189N are closely grouped within the High configuration. Interestingly, few mutations occur near the active site; only T219I in High and Q229R in Low are in this region. However, this static analysis alone does not provide a complete picture of the mutations' effects on overall protein stability or local structural changes, especially since many mutations are surface-exposed and thus may impact stability differently than those in more interior regions.

Additionally, a model obtained from Alphafill [16] for FNS I was analysed to represent the positioning of co-factors and substrates. Alphafill enhances Alphafold-predicted models by adding predicted co-factors and ligands based on similarities with homologous proteins. Although these models are approximations rather than exact replicas of the actual complexes, they offer a useful framework to estimate binding regions, such as for 2S-naringenin binding to the FNS I complexed enzyme.

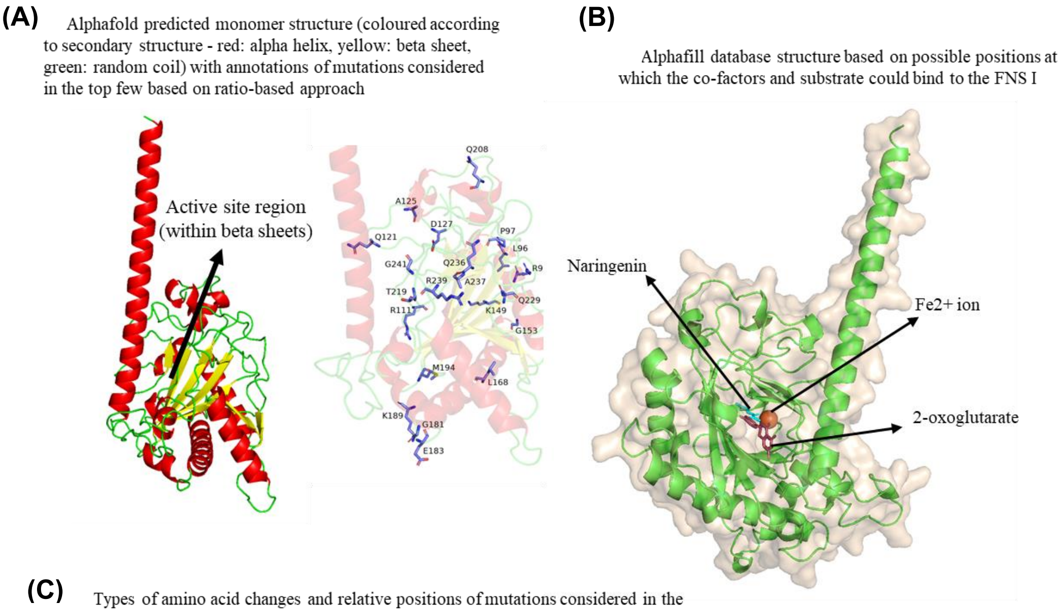

(C) Types of amino acid changes and relative positions of mutations considered in the top few based on ratio-based approach

| AA position | 91 | 96 | 97 | 111 | 121 | 125 | 127 | 149 | 153 | 168 |
|---|---|---|---|---|---|---|---|---|---|---|
| Location | Surface | Surface | Surface | Surface | Surface | Interior (random coil) | Interior (random coil) | Surface | Surface | Surface |
| WT<br>Type | R<br>+ | L<br>NPo | P<br>NPo | R<br>+ | Q<br>Po | A<br>NPo | D<br>- | K<br>+ | G<br>NPo | L<br>NPo |
| High<br>Type | | M<br>NPo | L<br>NPo | | E<br>- | | | T<br>Po | | I<br>NPo |
| Medium<br>Type | | | | | | V<br>NPo | E<br>- | | | |
| Low<br>Type | C<br>Npo | | | G<br>NPo | | | | | D<br>- | |

| AA position | 181 | 183 | 189 | 194 | 208 | 219 | 229 | 236 | 237 | 239 | 241 |
|---|---|---|---|---|---|---|---|---|---|---|---|
| Location | Surface | Surface | Surface | Surface | Surface | Near Active Site | Interior (beta sheet near active site) | Interior (beta sheet region) | Interior (beta sheet region) | Surface | Surface |
| WT<br>Type | G<br>NPo | E<br>- | K<br>+ | M<br>NPo | Q<br>Po | T<br>Po | Q<br>Po | Q<br>Po | A<br>NPo | R<br>+ | G<br>NPo |
| High<br>Type | V<br>NPo | K<br>+ | N<br>Po | V<br>NPo | | I<br>NPo | | | | | S<br>Po |
| Medium<br>Type | | | | | | | | | | | |
| Low<br>Type | | | | | H<br>+ | | R<br>+ | H<br>+ | S<br>Po | H<br>+ | |

| | |
|---|---|
| Npo | Non-polar |
| Po | Polar |
| + | Positively charged |
| - | Negatively charged |

Figure S21: Local effect of amino acid changes. (A) Putative structure of FNS I that predicted by AlphaFold, including the annotated mutation position designed using PUSDA. (B) Putative substrate and co-factors binding regions. (C) The type of local AA changes due to the specified substitutions in the FNS I variants that designed by PUSDA.

### Structural analysis of the PUSDA proposed enzymes

In the initial characterization of FNS I, potential dimerization was observed, as indicated by a second band approximately twice the size of the 48 kDa monomer in the 2D SDS-PAGE gel[2]. To explore this, AlphaFold 2[17] was used to predict the dimerization of two wild-type (WT)

models. The top five dimerization predictions highlighted possible mutation effects in configurations H1, H2, H3, L1, L2, and L3, based on whether the mutated amino acids are positioned near or away from dimerization regions, or within interior or surface regions.

AlphaFold predictions showed high confidence in the middle segment of the dimer model, suggesting this region may be reliably interpreted as having a functional impact on FNS I. For instance, in the highest-performing H3 configuration (E183K_K189N), residues E183 (negatively charged) and K189 (positively charged) are in proximity within a well-predicted region. This suggests that their combined mutation could affect enzyme function without significantly impacting dimerization, as these residues are distant from the dimerization interface. Changing K183 (positively charged) and N189 (polar, uncharged) could disrupt an ionic bond, possibly converting it to a weaker hydrogen bond, and potentially introduce an allosteric effect on FNS I function.

Similarly, the Q208 position appears in all L1, L2, and L3 configurations, while R91 and R111 are located within a region with a strong prediction score. Q208 (polar) lies on the protein surface, and changing it to H208 (positively charged) might influence the overall protein structure, potentially with a negative allosteric effect. Furthermore, R91 (positively charged, surface) changing to C91 (polar) and R111 (positively charged, near the dimerization interface) to G111 (neutral) could destabilize the protein and impact its interaction with reaction medium factors and dimerization potential.

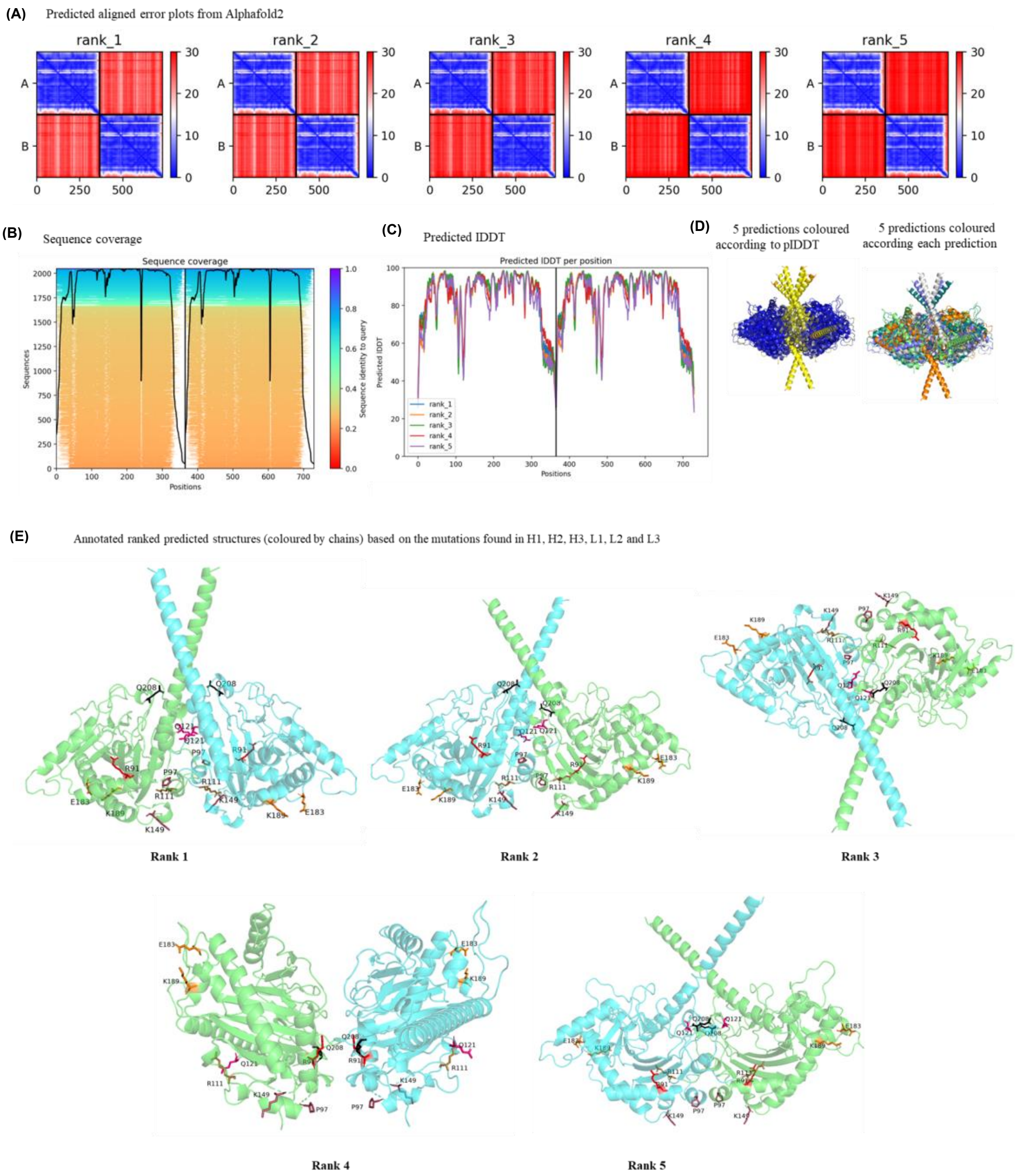

Figure S22: Structural analysis of the PUSDA proposed enzymes**.** (A) Prediction accuracy in 5 runs. (B) Sequence coverage. (C) Predicted IDDT. (D) Predicted structured with coloured according to pIDDT or each prediction. (E) Predicted structured and the location of the interested amino acids in each prediction

### Blind characterization of sorted 200 mutants

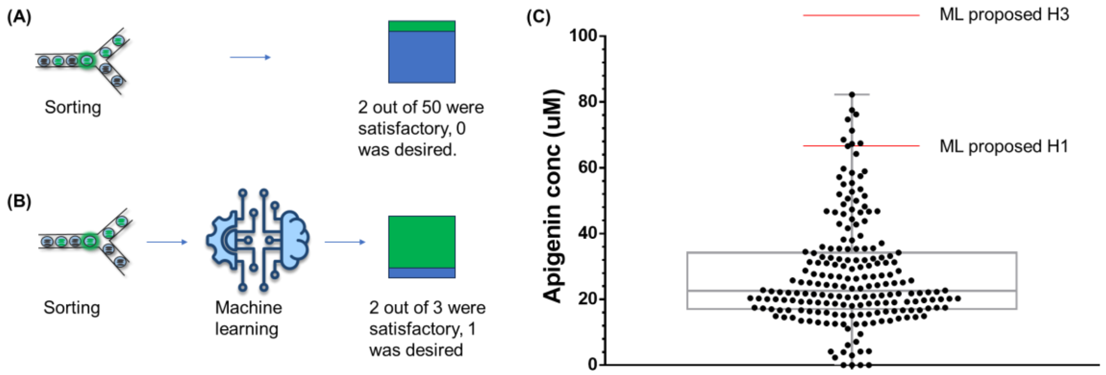

Figure S23: Efficiency of identifying high-performance has been significantly increased by using PUSDA. (A) Blindly characterizing the sorted mutants from H group. Only 4% of the sorted mutants achieved satisfactory productivity, and none of them achieved the desired productivity. (B) Mutants designed using ML method (PUDSA), resulting in 66.7% of the mutants being satisfactory, and one out of three achieved desired productivity. (C) Blind tested 200 mutants from the H group. 184 out of 200 (92%) showed improved apigenin production as compared to wild type, but only 8 out of the 200 (4%) had higher production than the ML model suggested H1. The highest production mutant from blind test is 48% lower than the ML model suggested H3 (production of 116.9 µM). Proving our ML assisted enzyme engineering framework significantly increased the efficiency of identification of high-performance enzymes as compared to previous technology.

### FNS mutation region

| Gene | Sequence | Note |
|---|---|---|
| FNS I | SEMTRLSREFFALPAEEKLEYDTTGGKRGGFTISTVL QGDDAMDWREFVTYFSYPINARDYSRWPKKPEGW RSTTEVYSEKLMVLGAKLLEVLSEAMGLEKGDLTK ACVDMEQKVLINYYPTCPQPDLTLGVRRHTDPGTIT ILLQDMVGGLQATRDGGKTWITVQ | Region of mutation |

Table 1: FNS I mutation region.
